# DNA methylation maintenance by DNMT1 is essential for human trophoblast stem cell homeostasis and differentiation

**DOI:** 10.64898/2026.07.02.735425

**Authors:** Sanam L Kavari, Yu Jin Jang, Gracie Clare Guerin, Laurel S Park, Elisia D Tichy, Joonhyuk Choi, Jonghwan Kim, Winifred Mak, Jennifer M Kalish

## Abstract

Proliferation of cytotrophoblasts (CTBs) and their differentiation into invasive extravillous trophoblasts (EVTs) are critical processes in early placental development. Defects in these processes are associated with adverse pregnancy outcomes, including recurrent pregnancy loss (RPL). There is evidence that reduced expression of the maintenance DNA methyltransferase DNMT1 in the placenta occurs in pregnancy loss and that RPL is associated with aberrant DNA methylation patterns. Therefore, we investigated the role of DNMT1 in human trophoblast growth and differentiation. Using human trophoblast stem cells (hTSCs), an *in vitro* analog to CTBs, we found that shRNA-mediated knockdown of *DNMT1* led to decreased hTSC proliferation, genome-wide reductions in methylation, broad changes in gene expression, and impaired EVT differentiation. Transcriptome profiling of *DNMT1*-deficient hTSCs and hTSC-derived EVTs highlighted aberrant cytokine expression, drawing a connection to prior reports of immunological dysfunction in RPL. Finally, using a catalytic DNMT1 chemical inhibitor, we demonstrate the canonical methyltransferase activity of DNMT1 is essential for EVT differentiation and invasion. This study identifies new roles for DNMT1 in trophoblasts and addresses the molecular basis of the associations between DNMT1 expression, altered DNA methylation profiles, and RPL.

## Introduction

The placenta is an essential organ for the establishment and maintenance of a healthy pregnancy. Defects in placental development and function are associated with adverse pregnancy outcomes, including preterm birth, intrauterine growth restriction, and both early and late pregnancy loss^1–3^. Early pregnancy loss, defined as intrauterine pregnancy without a viable fetus prior to 13 weeks gestation^4,5^, is among the most common complications, occurring in approximately 10% of all pregnancies^4,6,7^. Up to 5% of couples experience two or more pregnancy losses, meeting the definition for recurrent pregnancy loss (RPL)^8–10^. There are many possible etiologies of RPL, including genetic, structural, endocrine, and autoimmune factors; however, in more than half of cases, a definite cause of RPL is not identified^8,11,12^. Therefore, significant attention has been paid to identifying possible causes of pregnancy loss in patients with unexplained RPL.

Several studies have identified aberrant DNA methylation patterns in unexplained RPL placenta tissue^13–16^. Establishment and maintenance of DNA methylation is critical for cell lineage establishment in early embryonic development, so alterations to this process may disrupt the delicate balance of development and differentiation leading to pregnancy loss. DNA methylation is regulated by catalytic enzymes, known as DNA methyltransferases (DNMTs), that add methyl groups to the fifth carbon of cytosines (5mC) at predominantly CpG dinucleotides in development, differentiation, and cell division^17^. In mammals, three DNA methyltransferase enzymes, DNMT1, DNMT3A, and DNMT3B, are responsible for the establishment and maintenance of 5mC. DNMT3A and 3B add methylation *de novo* to DNA during development and differentiation^18,19^ while DNMT1 primarily maintains DNA methylation during cell division^20–23^. Reduced DNMT1 expression has been observed in the placental trophoblasts in early pregnancy loss^24^. However, the consequence of reduced DNMT1 abundance on human placental development has not been explored.

The human placenta is composed of many distinct cell types including the epithelial trophoblasts. The trophoblasts line the placental villi, interact directly with the maternal decidua, and perform many critical functions of the placenta^25^. The villous cytotrophoblasts (CTBs) are the proliferative and germinative trophoblast population^25,26^. The CTBs can further differentiate into syncytiotrophoblasts (STs), which produce hormones and participate directly in nutrient and gas exchange, or extravillous trophoblasts (EVTs), which invade into the maternal decidua and remodel the vasculature that supplies the placenta^25,27,28^. The establishment of *in vitro* human trophoblast stem cells (hTSCs), which are analogous to *in vivo* CTBs and can be differentiated into both STs and EVTs,^29^ has allowed for advances in the understanding of factors regulating trophoblast growth, self-renewal, and differentiation^30–33^. Recently, RPL patient-derived hTSCs have been established and demonstrate heterogeneous, cell line-specific defects in ST and EVT differentiation^34^. Additionally, it has been demonstrated that DNMT1 knockout (KO) is lethal in hTSCs and results in activation of previously silenced transposons^35^. While this work establishes that DNMT1 is essential in hTSCs, the KO approach was limited in its ability to assess the impact of longer depletion of DNMT1 and the role of DNMT1 in trophoblast differentiation.

Here we examine the role of DNMT1 in trophoblast self-renewal and differentiation using a knockdown (KD) approach in hTSCs. We established stable depletion of *DNMT1* for up to thirteen days in hTSC culture and characterized the impact of reduced *DNMT1* expression on hTSC self-renewal, growth, and methylation patterns. Moreover, we show an essential role for DNMT1 in EVT differentiation, which is dependent on its methyltransferase activity. Therefore, we establish a connection between reduced DNMT1 expression and trophoblast dysfunction in RPL.

## Results

### Knockdown of *DNMT1* disrupts hTSC morphology and proliferation

To investigate the role of DNMT1 in trophoblasts, we used a human trophoblast stem cell line (hTSC CT27) derived from first-trimester CTBs^29^. We reduced *DNMT1* expression in these hTSCs using lentivirus-based KD with two different shRNAs targeting *DNMT1* (shDNMT1_A and shDNMT1_B) compared to a control, non-targeting shRNA (shControl). Stable KD of *DNMT1* was confirmed by measuring transcript expression of *DNMT1* normalized to that of *GAPDH* by qPCR at days 3, 6, and 13 post-transduction (Fig. 1A, Fig. S1A). KD was also validated at the protein level on day 13 post-transduction by western blot (Fig. S1B & S1C). KD of *DNMT1* in hTSCs resulted in a flattening of the typical hTSC colony morphology (Fig. 1B). However, *DNMT1* KD did not alter the expression of genes associated with hTSC self-renewal, *GATA3, TP63,* and *TEAD4*, measured by qPCR (Fig. 1C), nor the expression of trophoblast markers TFAP2C and KRT7, measured by immunofluorescence (Fig. 1D & 1E). Despite the retained expression of self-renewal markers, *DNMT1* KD hTSCs display impaired proliferation as measured by EdU incorporation assay (Fig. 1F & 1G). These data show that expression of *DNMT1* is essential for maintaining the proliferative capacity of hTSCs, a hallmark of their stem cell identity.

**Figure 1:**
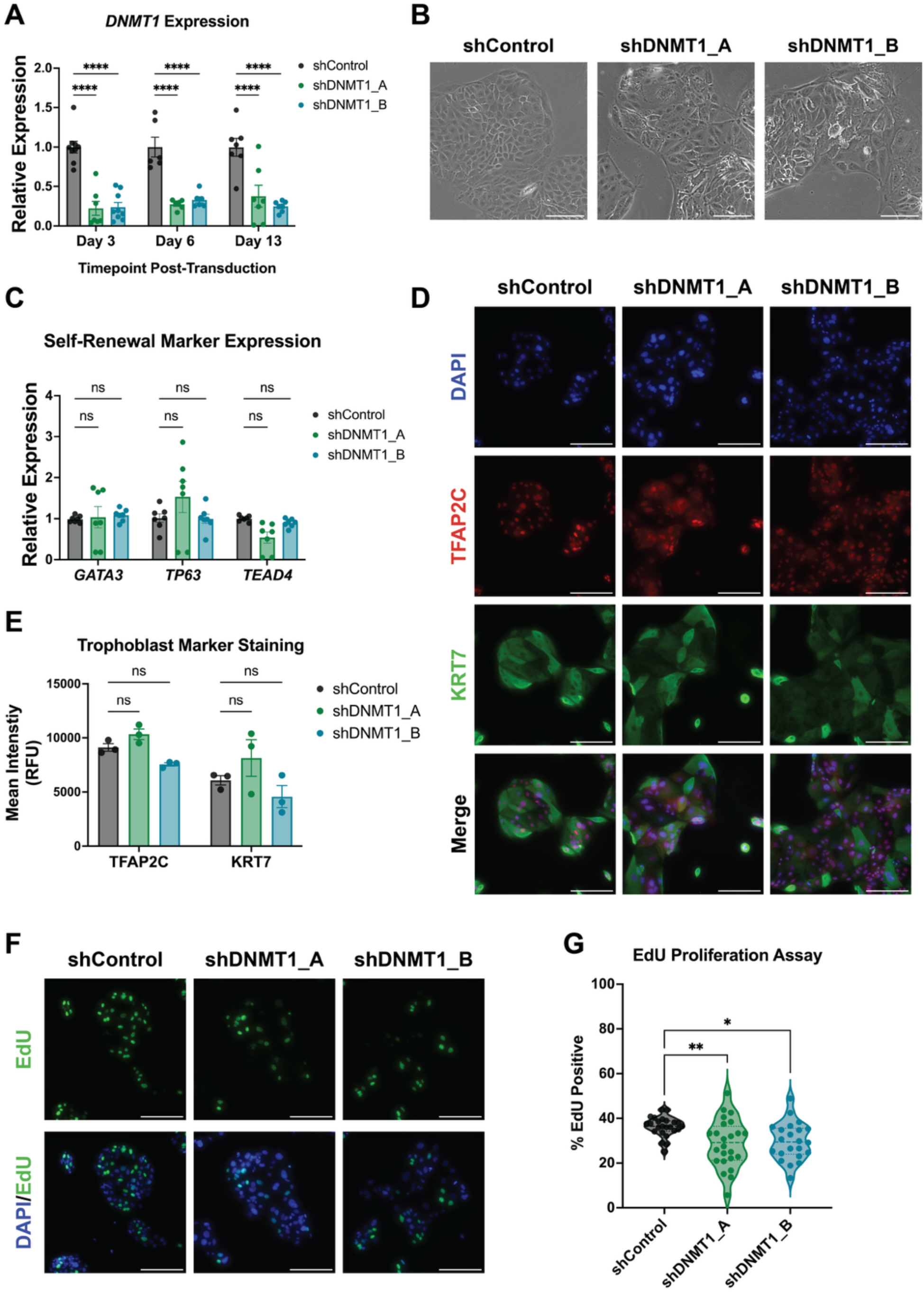
*DNMT1* KD in hTSCs alters morphology and impairs proliferation. A) Efficient KD of *DNMT1* at days 3, 6, and 13 post-transduction measured by qPCR normalized to *GAPDH* (Two-way ANOVA with multiple comparisons vs. shControl, n = 6-9 biological replicates). B) Representative phase-contrast images of *DNMT1* KD hTSCs compared to control hTSCs demonstrates altered morphology. C) *DNMT1* KD does not impact expression of markers associated with hTSC self-renewal *GATA3, TP63,* and *TEAD4* as measured by qPCR normalized to *GAPDH* (Two-way ANOVA with multiple comparisons vs. shControl, n = 7 biological replicates). D) Representative images of immunofluorescent staining of control and *DNMT1* KD hTSCs for trophoblast markers (Blue = DAPI, Red = TFAP2C, Green = KRT7). E) Quantification of trophoblast marker staining in demonstrates no change in expression between control and *DNMT1* KD hTSCs (Two-way ANOVA with multiple comparisons vs. shControl, n = 3 biological replicates, RFU = relative fluorescence units). F) Representative image of EdU proliferation assay shows reduced incorporation of EdU in *DNMT1* KD conditions compared to control (Blue = DAPI, Green = EdU) G) Quantification of EdU proliferation assay demonstrates reduced EdU positivity (One-way ANOVA with multiple comparisons vs. shControl, n = 22-24 images representing 5-6 biological replicates). (ns = not significant, * = p < 0.05, ** = p < 0.01, *** = p < 0.001, **** = p < 0.0001, scale bars represent 200 μm).

### Decreased *DNMT1* expression reduces global DNA methylation with differential impacts based on genomic context and chromatin state

To investigate the impact of reduced *DNMT1* expression on the methylome of hTSCs, we performed low-depth, whole genome bisulfite sequencing (Sparse-Seq) to measure global DNA methylation^36,37^ in hTSCs at day 13 post-transduction. Globally, *DNMT1* KD led to a reduction in DNA methylation at CpG dinucleotides from approximately 32% in shControl samples, a value consistent with previous observations^29,35,38^, to approximately 24% in both shDNMT1_A and shDNMT1_B samples (Fig. 2A, Data Table S1). This modest reduction in DNA methylation observed with *DNMT1* KD indicates that while DNMT1 does function to maintain DNA methylation in hTSCs, a portion of the hTSC methylome is likely actively established by the *de novo* DNA methyltransferases, DNMT3A and DNMT3B, as has been previously suggested^38^.

**Figure 2:**
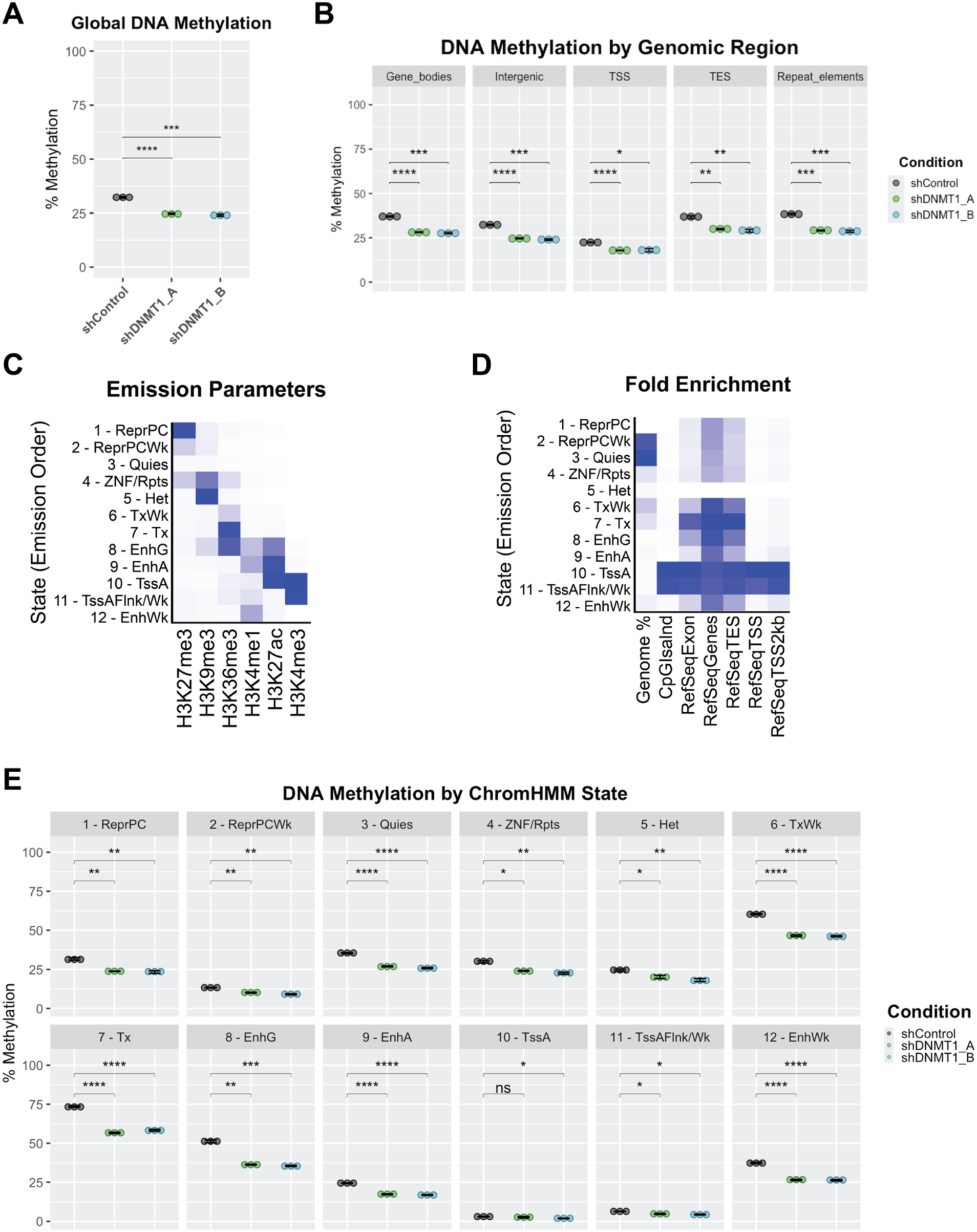
*DNMT1* KD in hTSC leads to global reductions in DNA methylation across genomic contexts. A) *DNMT1* KD results in a reduction in global CpG methylation as measured by Sparse-Seq. B) Stratification of Sparse-Seq data by genomic region demonstrates variable reductions in DNA methylation. C) ChromHMM state learning of chromatin enrichment data in hTSCs identified 12 chromatin states (ReprPC – Polycomb repressed, ReprPCWk – weakly Polycomb repressed, Quies – quiescent, ZNF/Rpts – zinc-finger protein genes and/or repeat elements, Het – constitutive heterochromatin, TxWk – weak transcription, Tx – transcription, EnhG – genic enhancers, EnhA – active enhancers, TssA – active transcription start sites, TssAFlnk/TssWk – regions flanking active transcription start sites or weak transcription start sites, EnhWk – weak enhancers). D) Validation of ChromHMM states by enrichment at various genomic contexts. E) Stratification of Sparse-Seq data by ChromHMM state reveals variation in the impact of *DNMT1* KD on DNA methylation by chromatin state. (T-tests with adjustments for multiple comparisons by Bonferroni method, ns = not significant, * = p < 0.05, ** = p < 0.01, *** = p < 0.001, **** = p < 0.0001, n = 3 biological replicates)

To further investigate the impacts of *DNMT1* KD on the hTSC methylome, we examined the methylation levels at defined genomic regions, namely in gene bodies, intergenic regions, transcription start sites (TSS) +/- 500bp, transcription end sites (TES) +/- 500bp, and repetitive elements, in *DNMT1* KD hTSCs. We found a significant reduction in DNA methylation across genomic compartments (Fig. 2B) in both shRNAs targeting *DNMT1*. However, TSS regions had the lowest baseline methylation levels in shControl hTSCs and the smallest magnitude of decreased methylation with *DNMT1* KD, from 22% in shControl hTSCs to 18% in shDNMT1 hTSCs (Data Table S1). The largest changes in DNA methylation were observed in gene bodies and repetitive elements, with an approximately 9% decrease in DNA methylation associated with *DNMT1* KD in both genomic contexts (Data Table S1).

To further understand the differential impacts of reduced *DNMT1* expression on DNA methylation in more granular genomic contexts, we performed ChromHMM^39,40^ chromatin state discovery analysis of publicly available ChIP-seq and CUT&TAG data in hTSCs. We used data for six histone marks, H3K27me3, H3K9me3, H3K36me3, H3K4me1, and H3K4me3,^30,38,41,42^ to define twelve chromatin states (Fig. 2C). These regions include states associated with repressive chromatin marks, H3K9me3 and H3K27me3, namely constitutive heterochromatin (Het) and regions strongly (ReprPC) and weakly (ReprPCWk) repressed by Polycomb. We also identified states associated with activating chromatin marks, H3K4me3, H3K4me1, and H3K27ac, including strong (Tx) and weak (TxWk) transcription, active enhancers (EnhA), weak or primed enhancers (EnhWk), genic enhancers (EnhG), active promoters (TssA), and regions flanking active promoters or weak promoters (TssAFlnk/Wk). We identified one state associated with mixed active (H3K36me3) and inactive (H3K9me3) chromatin marks, typically associated with zinc-finger protein genes and repetitive elements (ZNF/Rpts) and one quiescent (Quies) state associated with low levels of all chromatin marks (Fig. 2C)^39,40^. These states were validated by observing their overlap with defined genomic regions, including promoter associated states (TssA and TssAFlnk/Wk), which overlap most strongly with RefSeq defined TSSs and CpG Islands, transcription associated states (Tx and TxWk), which overlap most strongly with RefSeq defined genes, and the quiescent (Quies) state, which as expected accounts for the largest percent of the genome (Fig. 2D)^39,40^.

Once these states were defined, we stratified the Sparse-Seq DNA methylation data by ChromHMM state to investigate the impact of *DNMT1* KD on DNA methylation at these regions. We observed a significant reduction in DNA methylation at all chromatin states, apart from active promoters (TssA) (Fig. 2E, Data Table S1). Additionally, regions associated with promoters (TssA and TssAFlnk/Wk) were found to have the lowest baseline methylation (Fig. 2E). Regions associated with the highest levels of DNA methylation in shControl hTSCs are those associated with H3K36me3 (Tx, TxWk, and EnhG) (Data Table S1), consistent with prior reports of the overlap of high levels of DNA methylation and H3K36me3 in hTSCs^38^. The presence of H3K36me3 marks transcribed gene bodies and recruits the *de novo* DNA methyltransferases, DNMT3A and DNMT3B, to establish DNA methylation at these regions^43,44^. However, H3K36me3 marked regions are also associated with significant reductions in DNA methylation on *DNMT1* KD (Fig. 2E), indicating a role for the maintenance methyltransferase DNMT1 at these regions.

Previous studies have found that hTSCs have low levels of DNA methylation at placenta-specific partially methylated domains (PMDs)^29,38,45^. However, Toh *et al.* describe hTSC-specific PMDs which generally have lower levels of methylation compared to placental PMDs and are enriched for H3K27me3^45^. We observe intermediate levels of methylation at H3K27me3 marked regions (ReprPC and ReprPCWk) and a significant reduction in DNA methylation at these regions on depletion of *DNMT1* (Fig. 2E). This indicates a potential role for DNMT1 for maintaining DNA methylation at hTSC-specific PMDs. Other states with highly significant reductions in DNA methylation with *DNMT1* KD are unmarked, quiescent regions (Quies) and both active and weak enhancers (EnhA and EnhWk) (Fig. 2E). Taken together, these data show that DNMT1 functions primarily to maintain DNA methylation at transcribed gene bodies, repetitive regions, and regulatory elements in hTSCs.

### Knockdown of *DNMT1* impairs EVT differentiation but does not alter ST differentiation

To further assess the impact of *DNMT1* depletion on hTSC function, we investigated the effect of reduced *DNMT1* expression on hTSC differentiation capacity into both the EVT and ST lineages. After 4 days in selective media (6 days post-transduction), hTSCs transduced with either control or *DNMT1* targeting shRNAs were differentiated into hTSC-derived EVTs (hTSC-EVTs) or STs (hTSC-STs) following established protocols (Fig. S1)^29^. EVT and ST differentiation efficiency were measured after eight and six days of differentiation respectively.

Over the course of the eight-day EVT differentiation protocol, hTSCs acquire a distinctive, elongated mesenchymal morphology^29^. KD of *DNMT1* significantly alters EVT morphology with cells retaining the epithelial, colony-like morphology of undifferentiated hTSCs (Fig. 3A). Additionally, *DNMT1* KD EVTs display reduced expression of the characteristic EVT marker HLA-G as measured by immunofluorescence (Fig. 3B), qPCR (Fig. 3C), and flow cytometry (Fig. 3D & 3E). *DNMT1* KD hTSC-EVTs also demonstrate reduced expression of additional EVT markers *MMP2* and *ASCL2* compared to shControl hTSC-EVTs (Fig. 3C). These results indicate that reduced *DNMT1* expression impairs differentiation of hTSCs into EVTs *in vitro*.

**Figure 3:**
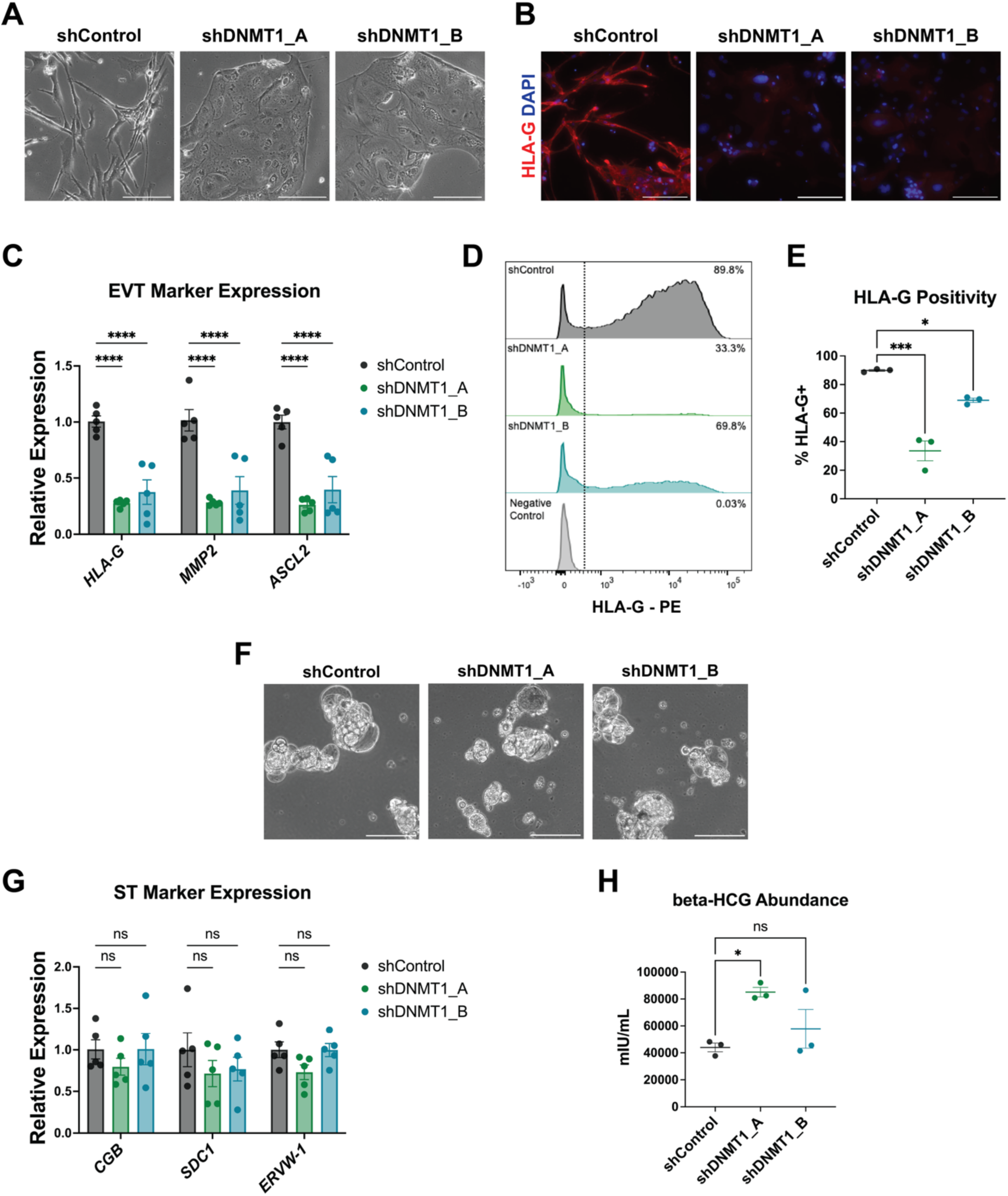
*DNMT1* KD impairs EVT differentiation but does not impact ST differentiation. A) Representative phase-contrast images show that *DNMT1* KD hTSC-EVTs fail to acquire the characteristic elongated, mesenchymal morphology associated with EVT differentiation as seen in shControl hTSC-EVTs. B) Immunofluorescent images show that *DNMT1* KD hTSC-EVTs do not express the EVT marker HLA-G at similar levels as shControl hTSC-EVTs (Blue = DAPI, Red = HLA-G). C) *DNMT1* KD results in a significant reduction in expression of EVT marker genes, *HLA-G, MMP2,* and *ASCL2*, as measured by qPCR normalized to *GAPDH* (Two-way ANOVA with multiple comparisons vs shControl, n = 5 biological replicates). D) Histogram of flow cytometry for HLA-G positive cells shows a reduction in HLA-G positivity in *DNMT1* KD hTSC-EVTs. E) Quantification of HLA-G positivity as measured by flow cytometry shows a significant reduction in HLA-G positive cells with *DNMT1* KD (One-way ANOVA with multiple comparisons vs shControl, n = 3 biological replicates). F) Representative phase-contrast images show that *DNMT1* KD does not impact the morphology of 3D hTSC-STs. G). *DNMT1* KD does not alter the expression of ST differentiation markers, *CGB, SDC1,* and *ERVW-1*, in hTSC-STs as measured by qPCR normalized to *GAPDH* (Two-way ANOVA with multiple comparisons vs shControl, n = 5 biological replicates). H) *DNMT1* KD also does not result in a consistent change in production of β-HCG as measured by ELISA on conditioned media (One-way ANOVA with multiple comparisons vs shControl, n = 3 biological replicates). (ns = not significant, * = p < 0.05, ** = p < 0.01, *** = p < 0.001, **** = p < 0.0001, scale bars represent 200 μm).

To examine the effects of *DNMT1* KD on the ST lineage, we differentiated hTSCs into hTSC-STs using a six-day, 3-dimensional differentiation protocol^29^. *DNMT1* KD does not affect the cystic morphology of differentiated hTSC-STs (Fig. 3F). Expression of ST markers, *CGB, SDC-1,* and *ERVW-1*, are also not affected by *DNMT1* KD (Fig. 3G). hTSC-STs mimic *in vivo* ST function, including production of hormones; therefore, we measured secretion of β-HCG into the medium by ELISA. We found β-HCG secretion was not consistently affected in hTSC-STs by *DNMT1* KD across the two shRNAs tested (Fig. 3H). These data show that reduced *DNMT1* expression does not significantly alter ST differentiation *in vitro*.

### Reduced *DNMT1* expression alters the hTSC and hTSC-EVT transcriptome leading to cytokine dysregulation

To further characterize the impact of *DNMT1* depletion on hTSCs and hTSC-EVTs, we employed whole-transcriptome analysis of control and *DNMT1* KD hTSCs and hTSC-EVTs by RNA-sequencing.

Principal components analysis (PCA) demonstrated that control and *DNMT1* KD undifferentiated hTSC samples clustered clearly with their replicates; however, clear clustering of the two DNMT1 shRNAs was not observed, potentially due to off-target effects specific to each shRNA (Fig. S2A). Differential gene expression analysis identified 1962 upregulated genes and 919 downregulated genes in shDNMT1_A hTSCs compared to shControl hTSCs (Fig. 4A, Data Table S2) and 608 upregulated genes and 212 downregulated genes in shDNMT1_B hTSCs compared to shControl hTSCs (Fig. 4B, Data Table S3).

**Figure 4:**
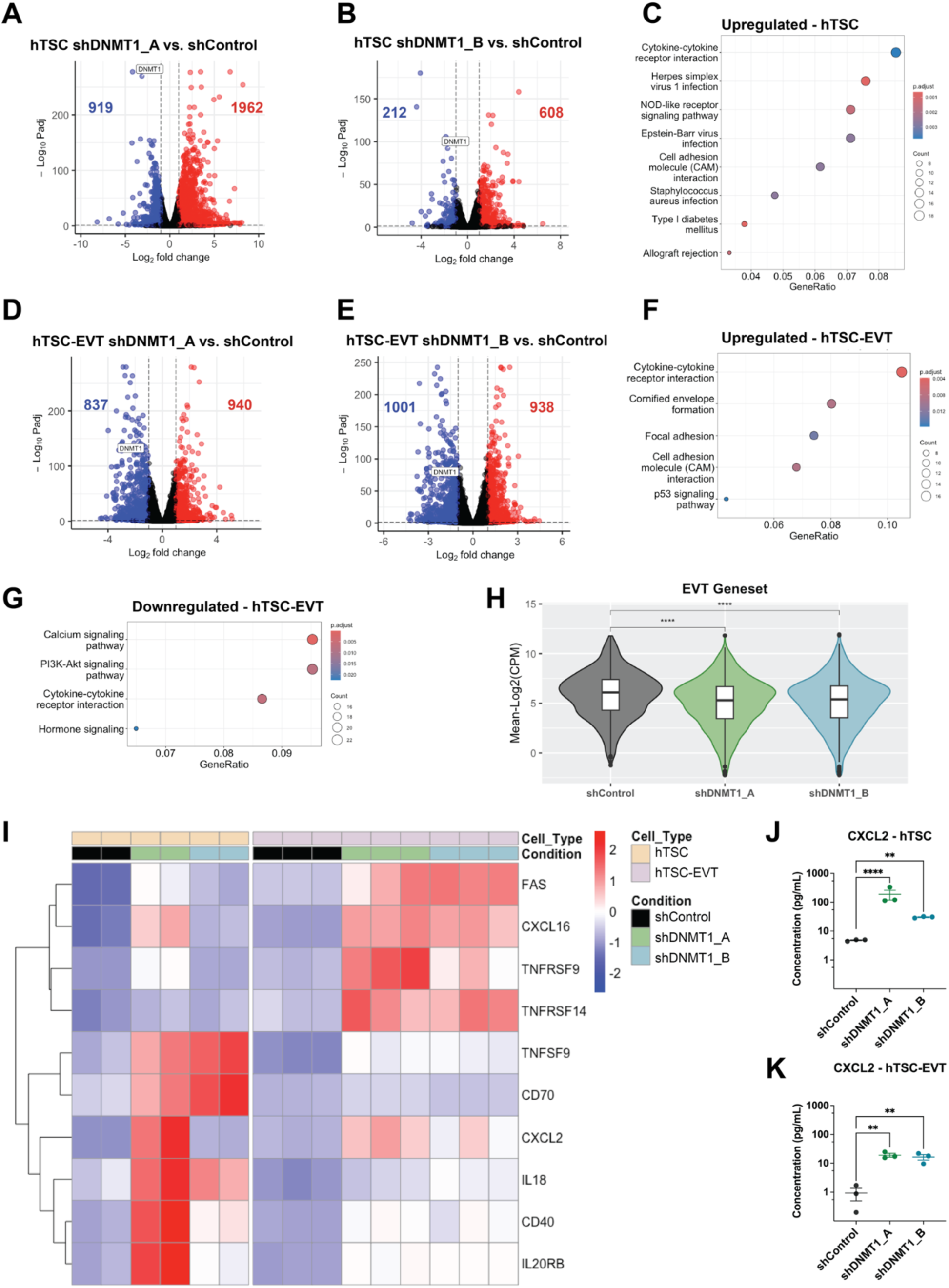
*DNMT1* KD impacts the hTSC and hTSC-EVT transcriptome leading to activation of cytokines and cytokine receptors. Volcano plots of A) shDNMT1_A vs shControl and B) shDNMT1_B vs shControl hTSC RNA-seq demonstrate gene dysregulation on depletion of *DNMT1* (padj > 0.05, |log_2_FoldChange| > 1, n = 2 biological replicates). C) KEGG term enrichment analysis of overlapping upregulated genes in shDNMT1_A and shDNMT1_B vs shControl comparisons reveals activation of immune associated pathways. Volcano plots of D) shDNMT1_A vs shControl and E) shDNMT1_B vs shControl hTSC-EVT RNA-seq show transcriptome level changes with KD of *DNMT1* (padj > 0.05, log_2_FoldChange > |1|, n = 3 biological replicates). KEGG term enrichment analysis of overlapping F) upregulated and G) downregulated genes in shDNMT1_A and shDNMT1_B vs shControl comparisons reveals dysregulation of pathways associated with immune dysfunction, interactions with the extracellular matrix, and pathways that include genes typically upregulated with EVT differentiation. H) Violin plots of EVT-associated gene set in control and *DNMT1* KD hTSC-EVTs shows failure to upregulate EVT-associated transcripts in shDNMT1 conditions (T-tests with adjustments for multiple comparisons by Bonferroni method). I) Heatmap of cytokine-cytokine reception pathway genes upregulated with *DNMT1* KD in hTSC and hTSC-EVT conditions highlights this pathway as the most highly dysregulated in both conditions. Measurement of CXCL2 production by ELISA on conditioned media in both J) hTSCs and K) hTSC-EVTs validates that upregulation of this gene at the RNA level in *DNMT1* KD correlates with increased protein production (One-way ANOVA with multiple comparisons vs shControl, n = 3 biological replicates). (** = p < 0.01, **** = p < 0.0001).

To identify genes whose dysregulation is attributable to reduced *DNMT1* expression, rather than shRNA specific off-target effects, we identified 472 overlapping upregulated genes and 107 overlapping downregulated genes in the shDNMT1_A vs shControl and shDNMT1_B vs. shControl comparisons (Fig. S2B & S2C, Data Table S4). Kyoto Encyclopedia of Genes and Genomes (KEGG) term enrichment analysis of these overlapping upregulated genes found that the most enriched class of genes was cytokines and their receptors (Fig. 4C), followed by gene sets associated with viral infections which are also driven primarily by cytokine dysregulation. Downregulated overlapping genes did not enrich for any KEGG terms; however, downregulated genes include positive cell cycle regulators, such as *CCNE2*^46^ and *CDC6*^47^, consistent with our finding that *DNMT1* KD impairs hTSC proliferation.

PCA of *DNMT1* KD hTSC-EVTs shows clear separation from shControl hTSC-EVTs on PC1, which accounted for 49.6% of total variance, showing a clear effect of *DNMT1* KD on the EVT transcriptome (Fig. S2D). Differential gene expression analysis identified 940 upregulated genes and 837 downregulated genes in the shDNMT1_A vs shControl comparison (Fig. 4D, Data Table S5) and 938 upregulated genes and 1001 downregulated genes in the shDNMT1_B vs. shControl comparison (Fig. 4E, Data Table S6). Between the two shDNMT1 vs. shControl conditions, we identified 420 overlapping upregulated genes and 574 overlapping downregulated genes (Fig. S2E & S2F, Data Table S7). Like the undifferentiated hTSC condition, KEGG term enrichment analysis of overlapping upregulated genes in *DNMT1* KD hTSC-EVTs also found the most enriched class of genes to be cytokines and their receptors (Fig. 4F). Downregulated genes were enriched for the PI3K/AKT and calcium signaling pathways, both of which include genes typically upregulated during EVT differentiation such as *ITGA1, FLT1, FLT4,* and *ERBB2* (Fig. 4G)^29,48^. Analysis of a defined gene set that is upregulated on EVT differentiation^29,48^ demonstrates a significant reduction in EVT associated transcripts in *DNMT1* KD hTSC-EVTs (Fig. 4H), indicating impaired EVT differentiation.

Due to the role of DNMT1 in maintaining methylation at imprinting control regions^49^, we investigated the expression of imprinted genes including placenta-specific imprinted genes^50^ in *DNMT1* KD hTSCs and hTSC-EVTs. We found four imprinted genes *SLC22A18*, *CPA4, SNRPN*, and *PEG10* were differentially expressed in *DNMT1* KD hTSCs and nine imprinted genes *CPA4*, *SGK2, CDKN1C, H19, ANO1, IGF2, LEP, MEG3,* and *MAGEL2* were differentially expressed in *DNMT1* KD hTSC-EVTs (Fig. S2G). However, the majority of the imprinted DEGs in hTSC-EVTs, namely *MEG3, SGK2, ANO1, IGF2, MAGEL2, CDKN1C, H19,* and *LEP* are not expressed in undifferentiated hTSCs or upregulated during EVT differentiation (Fig. S2G), so whether the change in expression of these genes is due to dysregulated imprinting or the observed impairment in EVT differentiation is not clear. Furthermore, the directionality of altered gene expression for many of the imprinted DEGs is not consistent with reduced methylation at their respective imprinting control region^51^, including *CPA4*^52^, *IGF2*^53^, *SLC22A18*^54^, *PEG10*^55^, and *SNRPN*^56^. Therefore, we do not observe changes in imprinted gene expression that are consistent with wide-spread imprinting disturbances in *DNMT1* KD hTSCs and hTSC-EVTs. This may be due to the multi-layered epigenetic regulation of imprinted gene expression that extends beyond DNA methylation.

Immune tolerance at the maternal-fetal interface is critical for the establishment of a healthy pregnancy^57^, so the upregulation of cytokines and their receptors in *DNMT1* KD hTSCs and hTSC-EVTs may offer insights into the mechanism by which reduced DNMT1 expression is associated with pregnancy loss^24^. To investigate this pathway further, we intersected the lists of upregulated genes in the cytokine-cytokine receptor interaction KEGG term in *DNMT1* KD hTSCs and hTSC-EVTs compared to control. We found 10 overlapping upregulated genes in this pathway in both *DNMT1* KD hTSCs and hTSC-EVTs (Fig. 4I). These include the proinflammatory cytokine *IL18*^58^, and both ligands and receptors in the tumor necrosis factor (TNF) superfamily, including *FAS, TNFRSF9, TNFRSF14, TNFSF9, CD70, CD40*, which function in T cell co-stimulation and cell killing^59^ (Fig. 4I). Furthermore, *CXCL2* and *CXCL1*6, members of the CXC chemokine family which function to regulate immune cell motility and recruitment^60^, are also upregulated in both *DNMT1* KD hTSCs and hTSC-EVTs (Fig. 4I). Expression of CXCL2 protein has previously been shown to be upregulated in the placental villi^61^ and immune cells in RPL^62,63^ and in a subset of RPL patient-derived hTSC lines^30^, highlighting this protein as a possible connection between reduced DNMT1 expression, impaired immune tolerance, and RPL. We validated that CXCL2 is secreted into the media at higher concentrations in both *DNMT1* KD hTSC and hTSC-EVTs compared to control by ELISA (Fig. 4J & 4K), demonstrating that upregulation of *CXCL2* at the RNA level corresponds to increased protein production. This analysis confirms that *DNMT1* KD impairs EVT differentiation and suggests aberrant expression of cytokines and their receptors as a possible mechanism by which reduced *DNMT1* expression impacts RPL.

### The DNMT1 catalytic methyltransferase activity is essential for EVT differentiation

Epigenetic enzymes often have catalytic and noncatalytic activities. To determine if the effect of *DNMT1* KD on EVT differentiation is directly related to its function as a DNA methyltransferase, we treated hTSCs with a catalytic DNMT1 inhibitor to assess the impact of the canonical function of DNMT1 in EVT differentiation. GSK3685032 (DNMT1i) functions as a non-covalent inhibitor of DNMT1 by reversibly binding to the catalytic, active site and does not robustly affect DNMT1 protein levels^64^. We treated hTSCs with 10 μM DNMT1i or DMSO as a vehicle control for 72 hours, to mimic the timeline of DNMT1 depletion in the *DNMT1* KD experiments and subsequently followed the eight-day EVT differentiation protocol with continued DNMT1i treatment. At the end of the EVT differentiation protocol, we confirmed that treatment with DNMT1i did not reduce DNMT1 protein levels by western blot (Fig. 5A & 5B). Furthermore, we validated that treatment with DNMT1i led to reduced methylation at the *KCNQ1OT1*:TSS-DMR (Fig. 5C), an imprinting control region whose methylation is maintained by DNMT1^49^, by targeted bisulfite sequencing of the region. This demonstrates that treatment with DNMT1i impairs the DNA methylation maintenance function of DNMT1 without influencing protein abundance.

**Figure 5:**
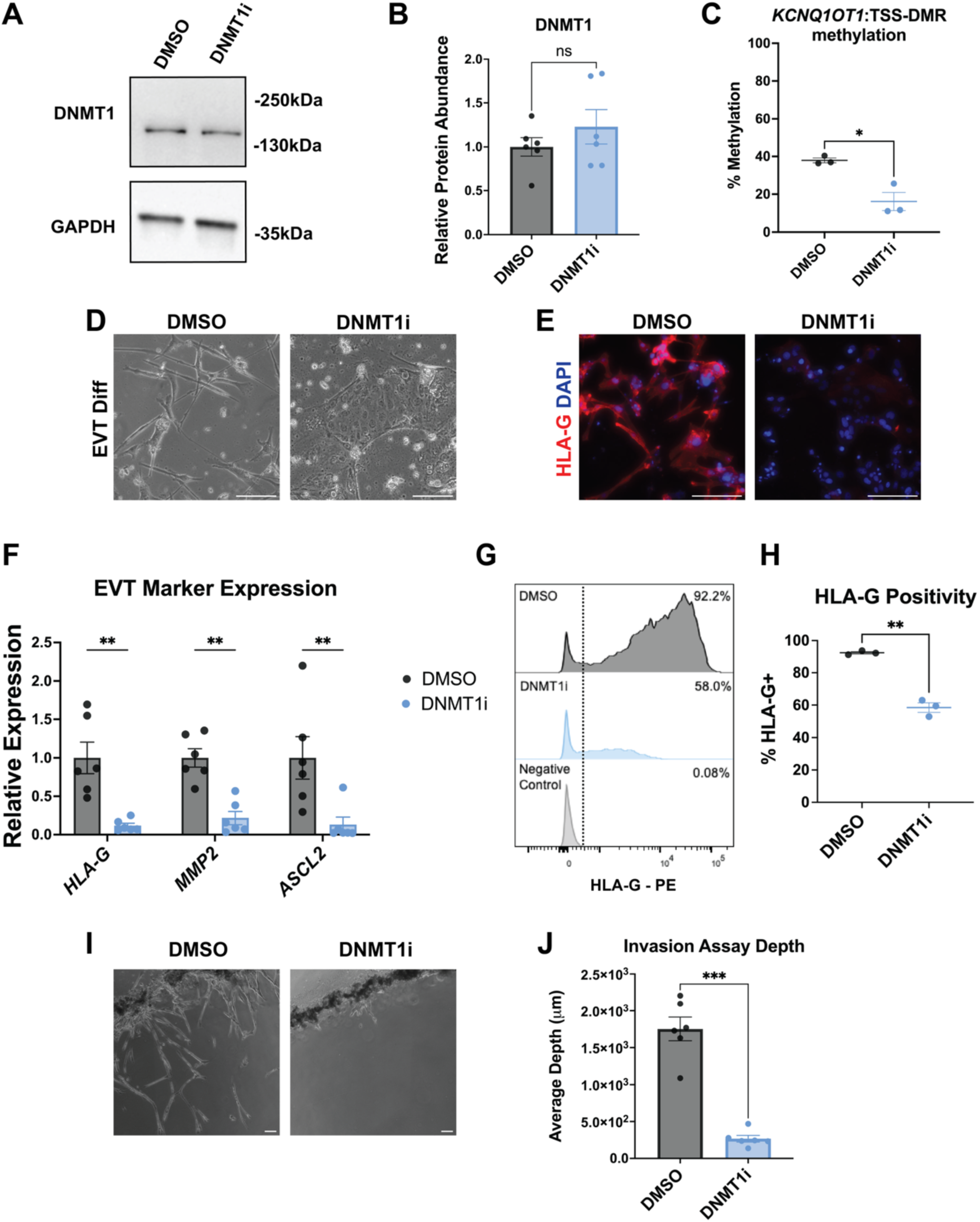
Catalytic activity of DNMT1 is essential for hTSC-EVT differentiation. A) Representative western blot for DNMT1 with GAPDH as a loading control on day 8 of EVT differentiation demonstrates that catalytic DNMT1 inhibitor (DNMT1i) does not reduce DNMT1 protein abundance compared to DMSO treated cells. B) Quantification of western blots demonstrating retained DNMT1 protein abundance normalized to GAPDH in DNMT1i compared to DMSO (Welch’s t-test, n = 6 biological replicates). C) Targeted bisulfite sequencing of the *KCNQ1OT1*:TSS-DMR region demonstrates reduction in methylation with treatment with DNMT1i indicating inhibition of DNMT1 methyltransferase activity (Welch’s t-test, n = 3 biological replicates). D) Representative phase-contrast images of DNMT1i treated hTSC-EVTs compared to DMSO treated cells as a control shows failure to take on characteristic *in vitro* EVT morphology. B) Immunofluorescent images show that DNMT1i treated hTSC-EVTs do not express the EVT marker HLA-G at similar levels as control DMSO treated hTSC-EVTs (Blue = DAPI, Red = HLA-G). F) qPCR shows a significant reduction in expression of EVT marker genes, *HLA-G, MMP2,* and *ASCL2*, with DNMT1i treatment compared to DMSO (Two-way ANOVA with multiple comparisons, n = 6 biological replicates). G) Histogram of flow cytometry for HLA-G shows a reduction in HLA-G positivity with treatment with DNMT1i. H) Quantification of HLA-G positivity as measured by flow cytometry shows a significant reduction in HLA-G positive cells with DNMT1i treatment (Welch’s t-test, n = 3 biological replicates). I) Representative phase-contrast images of Matrigel invasion assay shows impaired invasive function of DNMT1i treated hTSC-EVTs. J) Quantification of Matrigel invasion assay shows a significant reduction in invasion depth with DNMT1i treatment (Welch’s t-test, n = 6 biological replicates). (ns = not significant, * = p < 0.05, ** = p < 0.01, *** = p < 0.001, scale bars represent 200 μm).

We next aimed to assess the impact of DNMT1i treatment on EVT differentiation efficiency of hTSCs. We found treatment with DNMT1i significantly altered EVT morphology (Fig. 5D), with retention of the epithelial morphology seen in hTSCs prior to EVT differentiation, as observed with *DNMT1* KD (Fig. 3A). Additionally, DNMT1i treatment significantly reduced HLA-G expression as measured by immunofluorescence (Fig. 5E), qPCR (Fig. 5F), and flow cytometry (Fig 5G & 5H), indicating impaired EVT differentiation. Treatment with DNMT1i also led to reduced expression of two additional EVT markers, *MMP2* and *ASCL2* (Fig. 5F). The reductions in EVT marker expression on treatment with DNMT1i were of greater magnitude than the reductions seen with *DNMT1* KD (Fig. 3C), potentially indicating more complete inhibition of DNMT1 with the chemical inhibitor treatment compared to the partial depletion with the KD approach. Finally, because the critical function of EVTs is to invade into the maternal decidua, we investigated the impact of DNMT1 inhibition on hTSC-EVT invasion. We found that DNMT1i treatment significantly reduced EVT invasion depth (Fig. 5I & 5J). Therefore, inhibition of the catalytic function of DNMT1 is sufficient to impair EVT differentiation and function in the absence of reduced protein abundance, indicating that the DNA methyltransferase function of DNMT1 is essential for EVT differentiation.

## Discussion

In this report, we aimed to elucidate the functional consequence of reduced DNMT1 protein expression in the placental villi in pregnancy loss^24^ by using hTSCs as an *in vitro* trophoblast model. We demonstrate that shRNA-mediated KD of *DNMT1* results in reduced proliferative capacity, genome-wide reductions in DNA methylation, broad gene expression changes, and impaired EVT differentiation. Prior work using a CRISPR KO approach demonstrated that expression of *DNMT1* is essential in hTSCs, with cultures of KO cells being overgrown by un-edited cells within ten days^35^. While this previous work did not identify a reduction in proliferation with *DNMT1* KO^35^, this difference may be due to their more acute, complete depletion of *DNMT1* compared to our longer, partial depletion. Furthermore, our study mimics the diminished level of DNMT1 protein found in villous CTBs of early pregnancy loss, providing crucial insight into human miscarriage, an *in vivo* process which has been difficult to study due to ethical constraints. Interestingly, although we found that reduced *DNMT1* expression impaired proliferation, a hallmark of stem cell identity, this occurred in the absence of changes in expression of hTSC self-renewal genes.

Our finding that DNMT1 depletion impairs hTSC proliferation and the finding that DNMT1 is essential in hTSCs^35^ contrast prior work in mTSCs, where knockout of DNMT1 along with DNMT3A and DNMT3B do not impact self-renewal^65^. A similar pattern has been observed in hESCs, where DNMT1 knockout leads to rapid cell death^66^, and mESCs, where DNMT1 knockout is tolerated^22^. The dependence of hESCs on DNMT1 has been attributed in part to the different pluripotent state represented by hESCs compared to mESCs^66^. hTSCs and mTSCs may also represent distinct cell states as trophoblast lineage establishment is significantly different between human and mouse. CRISPR screening of hTSCs found that most transcription factors that are essential for maintaining mTSC self-renewal are dispensable for hTSCs, while transcription factors essential for the development of more terminally differentiated mouse trophoblast lineages significantly influenced hTSC growth and self-renewal^31^. Therefore, hTSCs may represent a more differentiated stem cell population than mTSCs and a state in which DNMT1 activity is essential.

Our characterization of the methylation patterns in hTSCs is also consistent with prior reports^29,35,38,45^. Similar to Lea *et al*., we identify the highest levels of DNA methylation in regions overlapping H3K36me3 marked transcribed gene bodies^38^. We also find depletion of DNA methylation with *DNMT1* KD at repetitive elements, consistent with described role of DNMT1 in silencing transposons in hTSCs^35^. Additionally, prior reports have also identified a general depletion of DNA methylation at placenta-specific PMDs in hTSCs but instead identify hTSC-specific PMDs^29,38,45^. Placental-PMDs tend to be enriched for H3K9me3, indicating their association with heterochromatic domains, while hTSC-PMDs are more associated with Polycomb repressed H3K27me3 regions^45^. This is consistent with our finding of intermediate methylation levels at regions enriched for H3K27me3 (ReprPC and ReprPCWk), We also found a reduction in methylation at these H3K27me3 regions on *DNMT1* KD, indicating a role for DNMT1 in maintaining hTSC-PMDs. DNMT1 also functions to maintain DNA methylation at imprinting control regions^49^. While we could not assess methylation at specific imprinting control regions due to limitations of the low-depth bisulfite sequencing, our analysis of imprinted gene expression indicates that imprinting disturbances are not likely to underlie the role of DNMT1 in trophoblast self-renewal and differentiation. Therefore, using the recently developed Sparse-Seq approach^36^, we successfully recapitulate previously reported hTSC methylation profiles generated with higher-depth approaches and demonstrate the impact of *DNMT1* depletion on broad genomic regions.

A further consequence of *DNMT1* depletion on hTSC function is an impairment of EVT differentiation potential, without changes in potential for ST differentiation. DNMT1 has been shown to play a role in lineage specification in a variety of contexts^67–72^, and the process of EVT differentiation requires dynamic epigenetic transitions including changes in DNA methylation^30,73,74^. Overexpression of *DNMT3L*, a co-factor of the *de novo* DNA methyltransferases DNMT3A and DNMT3B, was recently shown to impair differentiation of hTSCs into the ST lineage^38^. This emphasizes the importance of balanced DNA methylation for trophoblast differentiation toward both terminally differentiated lineages. Using a catalytic, chemical inhibitor of DNMT1 we demonstrate that the EVT differentiation defect we observed is dependent on the methyltransferase activity of DNMT1 and that reduced DNMT1 activity impairs not only the differentiation but also the invasive function of EVTs. Impaired trophoblast differentiation and invasion provide a functional link between reduced expression of DNMT1 in the trophoblast^24^, alterations in placental methylation patterns^13–15^, and pregnancy loss.

Finally, we assessed the transcriptome of *DNMT1-*deficient hTSC-EVTs. We found that reduced *DNMT1* expression resulted in aberrant expression of cytokines and their receptors in both hTSCs and hTSC-EVTs. The establishment and maintenance of immune tolerance at the maternal-fetal interface is a critical function of EVTs; however, the role of immunological dysfunction in RPL is controversial^57^. Significant changes in immune cell profiles and cytokine expression have been observed in RPL tissue^62,63,75,76^ and in hTSCs derived from RPL pregnancies^34^, but if these changes precede pregnancy loss or are a consequence of pregnancy loss has been difficult to address. We cannot directly address temporality in our *in vitro* study, but our work does provide a compelling connection between alterations in DNMT1 expression and DNA methylation in pregnancy loss^13–15,24^ and cytokine dysregulation in the background of hTSCs derived from normal pregnancy.

Furthermore, while we do not directly establish if cytokine dysregulation contributes to the EVT differentiation defect observed on *DNMT1* depletion, other studies have reported direct effects of cytokines on EVT differentiation efficiency and function^77,78^. Therefore, determining if the cytokines we observed to be upregulated on *DNMT1* depletion have direct impacts on trophoblast differentiation offers a pathway to further interrogate molecular mechanisms underlying RPL. The cytokines we observed to be upregulated on *DNMT1* depletion represent a group of proinflammatory cytokines involved in immune cell chemotaxis, including *CXCL2.* We validated CXCL2 protein abundance was also increased in *DNMT1* KD hTSCs and hTSC-EVTs by ELISA. CXCL2 protein has previously been observed to be upregulated in the placenta in RPL and in *in vitro* trophoblasts in response to depletion of UHRF1, a binding partner of DNMT1^61^. Therefore, DNMT1 and its binding partner, UHRF1, may work in concert to maintain appropriate expression of immune-modulating factors, like *CXCL2*, at the maternal-fetal interface.

Taken together, we describe novel roles of DNMT1 in human trophoblasts and provide insights into molecular underpinnings of RPL. Our findings add to recent work demonstrating the role of epigenetic modifiers in trophoblast growth and differentiation^35,38,79–82^. This growing body of literature emphasizes the unique epigenetic mechanisms underlying cell growth and differentiation in the extraembryonic lineage.

## Materials and Methods

### Key Resources Table

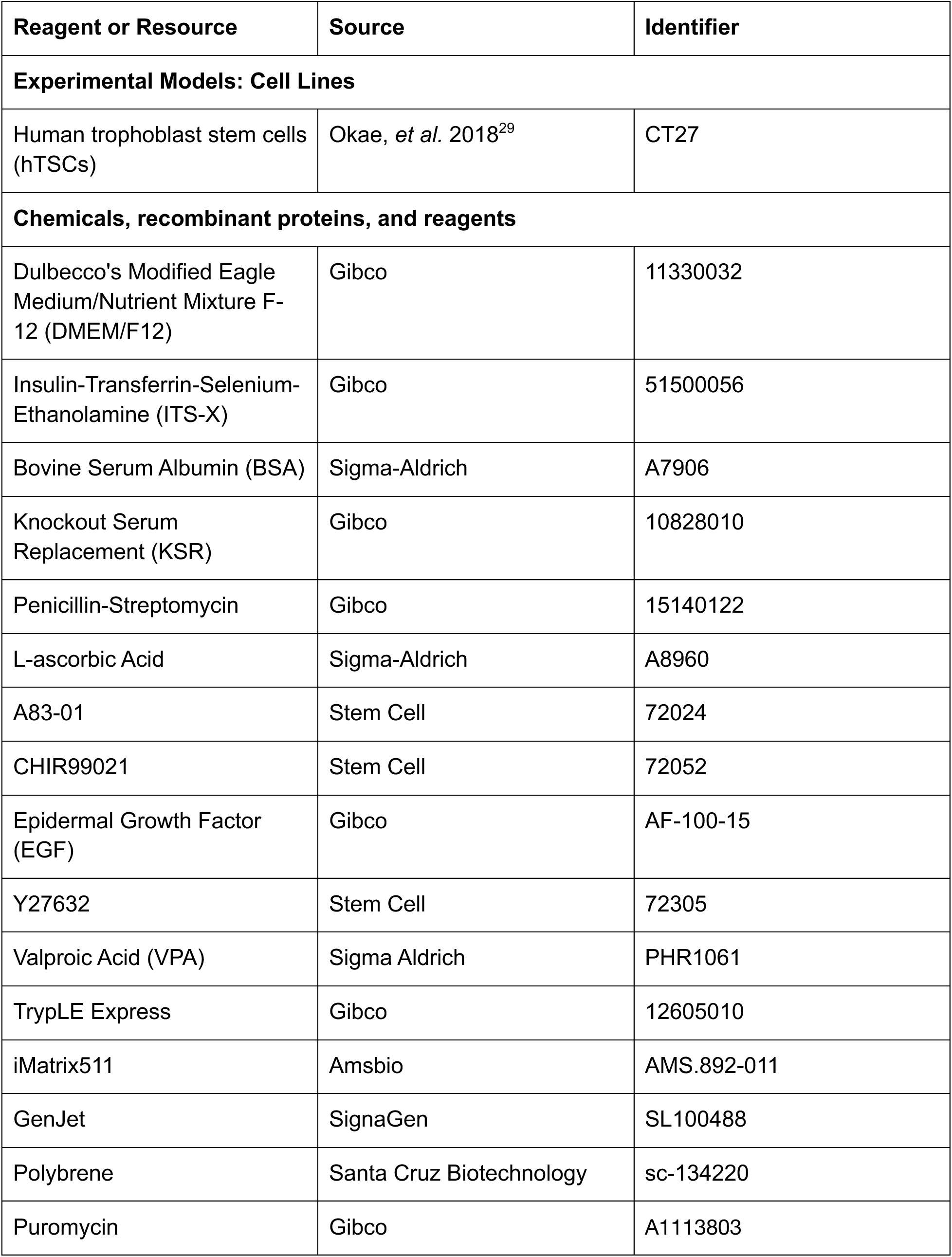
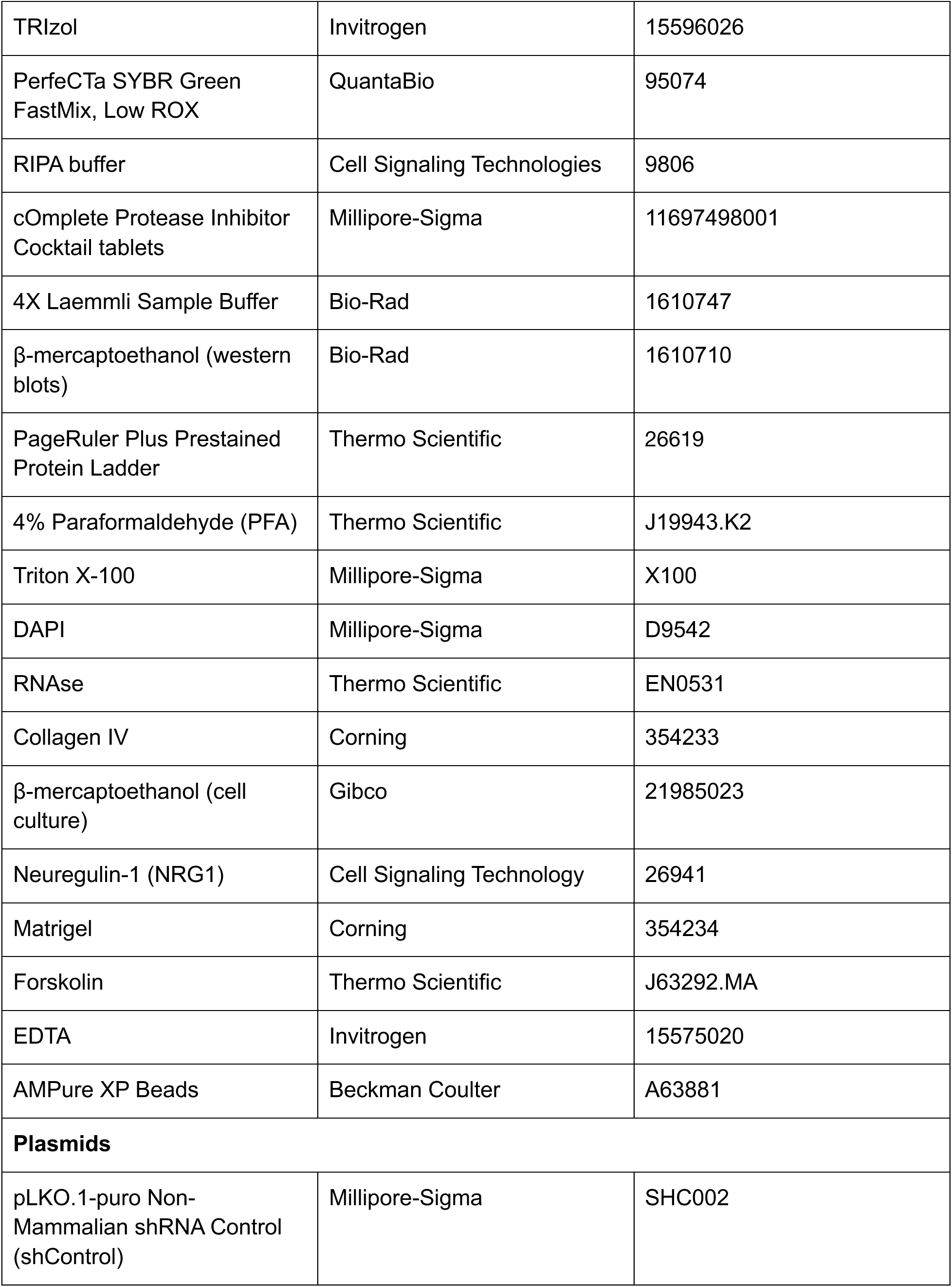
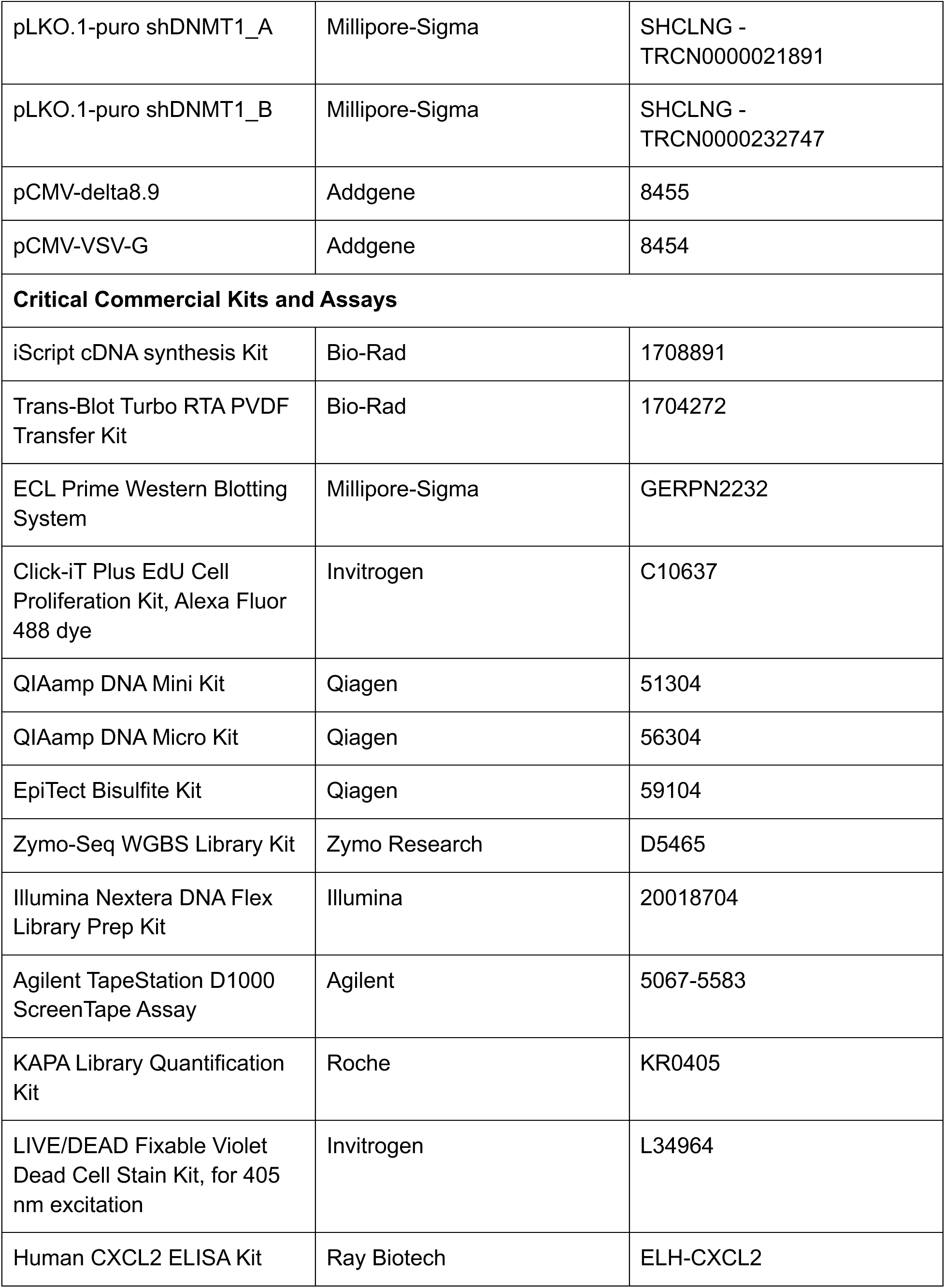
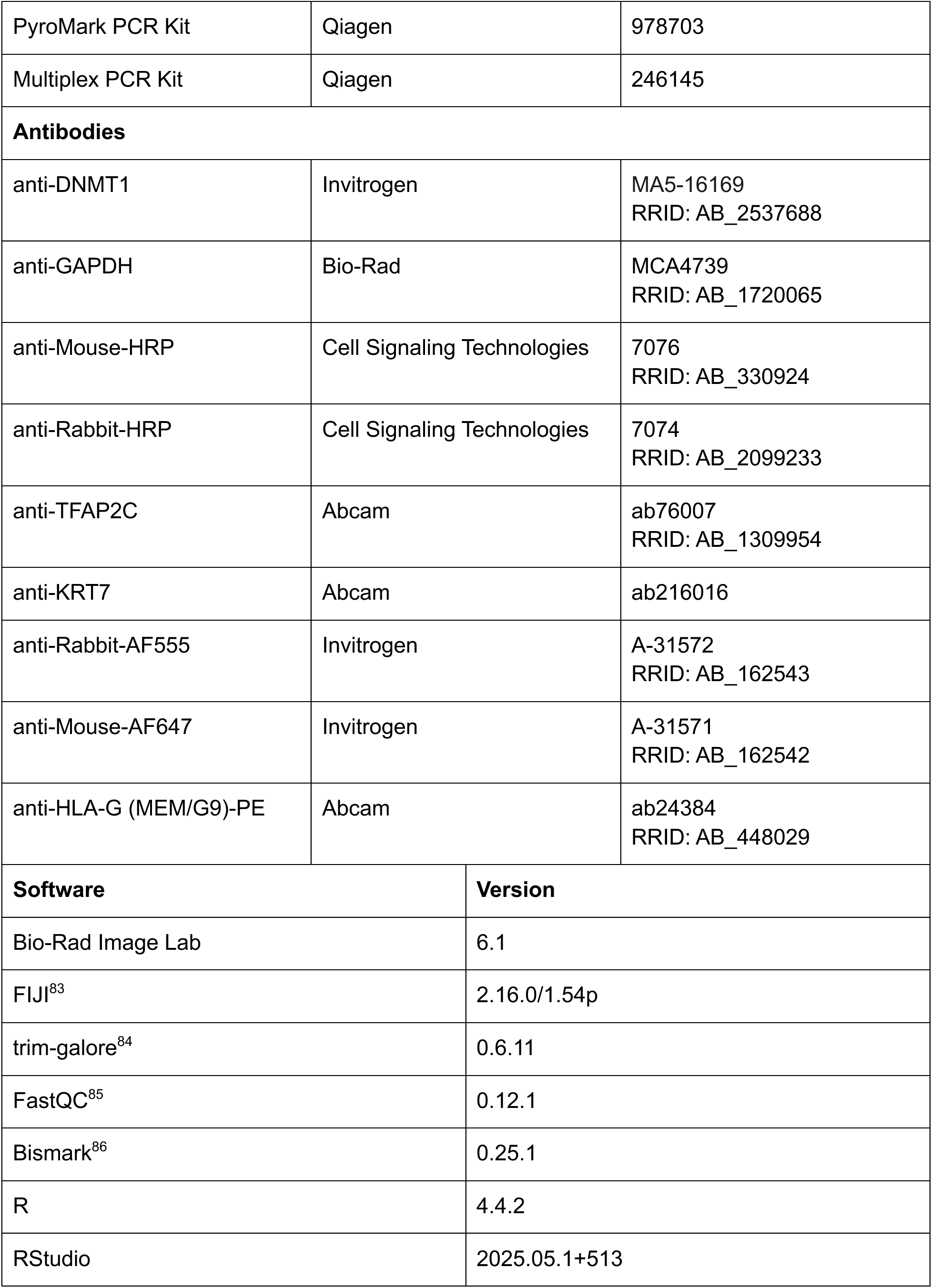
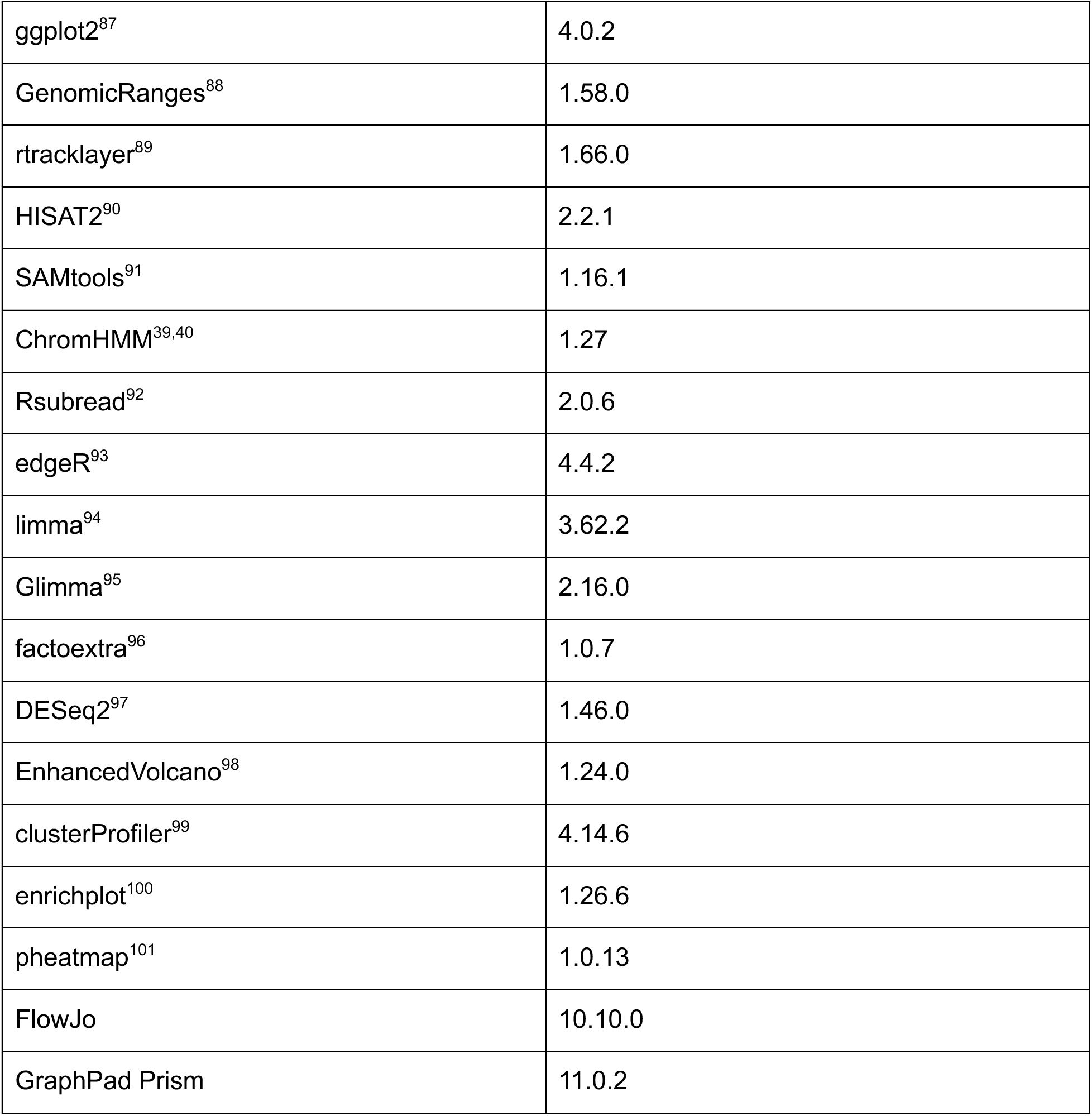

| Reagent or Resource | Source | Identifier |
| --- | --- | --- |
| **Experimental Models: Cell Lines** |  |  |
| Human trophoblast stem cells (hTSCs) | Okae, *et al.* 2018<sup>29</sup> | CT27 |
| **Chemicals, recombinant proteins, and reagents** |  |  |
| Dulbecco's Modified Eagle Medium/Nutrient Mixture F-12 (DMEM/F12) | Gibco | 11330032 |
| Insulin-Transferrin-Selenium-Ethanolamine (ITS-X) | Gibco | 51500056 |
| Bovine Serum Albumin (BSA) | Sigma-Aldrich | A7906 |
| Knockout Serum Replacement (KSR) | Gibco | 10828010 |
| Penicillin-Streptomycin | Gibco | 15140122 |
| L-ascorbic Acid | Sigma-Aldrich | A8960 |
| A83-01 | Stem Cell | 72024 |
| CHIR99021 | Stem Cell | 72052 |
| Epidermal Growth Factor (EGF) | Gibco | AF-100-15 |
| Y27632 | Stem Cell | 72305 |
| Valproic Acid (VPA) | Sigma Aldrich | PHR1061 |
| TrypLE Express | Gibco | 12605010 |
| iMatrix511 | Amsbio | AMS.892-011 |
| GenJet | SignaGen | SL100488 |
| Polybrene | Santa Cruz Biotechnology | sc-134220 |
| Puromycin | Gibco | A1113803 |
| TRIzol | Invitrogen | 15596026 |
| PerfeCTa SYBR Green FastMix, Low ROX | QuantaBio | 95074 |
| RIPA buffer | Cell Signaling Technologies | 9806 |
| cOmplete Protease Inhibitor Cocktail tablets | Millipore-Sigma | 11697498001 |
| 4X Laemmli Sample Buffer | Bio-Rad | 1610747 |
| β-mercaptoethanol (western blots) | Bio-Rad | 1610710 |
| PageRuler Plus Prestained Protein Ladder | Thermo Scientific | 26619 |
| 4% Paraformaldehyde (PFA) | Thermo Scientific | J19943.K2 |
| Triton X-100 | Millipore-Sigma | X100 |
| DAPI | Millipore-Sigma | D9542 |
| RNAse | Thermo Scientific | EN0531 |
| Collagen IV | Corning | 354233 |
| β-mercaptoethanol (cell culture) | Gibco | 21985023 |
| Neuregulin-1 (NRG1) | Cell Signaling Technology | 26941 |
| Matrigel | Corning | 354234 |
| Forskolin | Thermo Scientific | J63292.MA |
| EDTA | Invitrogen | 15575020 |
| AMPure XP Beads | Beckman Coulter | A63881 |
| **Plasmids** |  |  |
| pLKO.1-puro Non-Mammalian shRNA Control (shControl) | Millipore-Sigma | SHC002 |
| pLKO.1-puro shDNMT1_A | Millipore-Sigma | SHCLNG -<br>TRCN0000021891 |
| pLKO.1-puro shDNMT1_B | Millipore-Sigma | SHCLNG -<br>TRCN0000232747 |
| pCMV-delta8.9 | Addgene | 8455 |
| pCMV-VSV-G | Addgene | 8454 |
| **Critical Commercial Kits and Assays** |  |  |
| iScript cDNA synthesis Kit | Bio-Rad | 1708891 |
| Trans-Blot Turbo RTA PVDF<br>Transfer Kit | Bio-Rad | 1704272 |
| ECL Prime Western Blotting<br>System | Millipore-Sigma | GERPN2232 |
| Click-iT Plus EdU Cell<br>Proliferation Kit, Alexa Fluor<br>488 dye | Invitrogen | C10637 |
| QIAamp DNA Mini Kit | Qiagen | 51304 |
| QIAamp DNA Micro Kit | Qiagen | 56304 |
| EpiTect Bisulfite Kit | Qiagen | 59104 |
| Zymo-Seq WGBS Library Kit | Zymo Research | D5465 |
| Illumina Nextera DNA Flex<br>Library Prep Kit | Illumina | 20018704 |
| Agilent TapeStation D1000<br>ScreenTape Assay | Agilent | 5067-5583 |
| KAPA Library Quantification<br>Kit | Roche | KR0405 |
| LIVE/DEAD Fixable Violet<br>Dead Cell Stain Kit, for 405<br>nm excitation | Invitrogen | L34964 |
| Human CXCL2 ELISA Kit | Ray Biotech | ELH-CXCL2 |
| PyroMark PCR Kit | Qiagen | 978703 |
| Multiplex PCR Kit | Qiagen | 246145 |
| **Antibodies** |  |  |
| anti-DNMT1 | Invitrogen | MA5-16169<br>RRID: AB_2537688 |
| anti-GAPDH | Bio-Rad | MCA4739<br>RRID: AB_1720065 |
| anti-Mouse-HRP | Cell Signaling Technologies | 7076<br>RRID: AB_330924 |
| anti-Rabbit-HRP | Cell Signaling Technologies | 7074<br>RRID: AB_2099233 |
| anti-TFAP2C | Abcam | ab76007<br>RRID: AB_1309954 |
| anti-KRT7 | Abcam | ab216016 |
| anti-Rabbit-AF555 | Invitrogen | A-31572<br>RRID: AB_162543 |
| anti-Mouse-AF647 | Invitrogen | A-31571<br>RRID: AB_162542 |
| anti-HLA-G (MEM/G9)-PE | Abcam | ab24384<br>RRID: AB_448029 |
| **Software** |  | **Version** |
| Bio-Rad Image Lab |  | 6.1 |
| FIJI<sup>83</sup> |  | 2.16.0/1.54p |
| trim-galore<sup>84</sup> |  | 0.6.11 |
| FastQC<sup>85</sup> |  | 0.12.1 |
| Bismark<sup>86</sup> |  | 0.25.1 |
| R |  | 4.4.2 |
| RStudio |  | 2025.05.1+513 |

|  |  |
| --- | --- |
| ggplot2<sup>87</sup> | 4.0.2 |
| GenomicRanges<sup>88</sup> | 1.58.0 |
| rtracklayer<sup>89</sup> | 1.66.0 |
| HISAT2<sup>90</sup> | 2.2.1 |
| SAMtools<sup>91</sup> | 1.16.1 |
| ChromHMM<sup>39,40</sup> | 1.27 |
| Rsubread<sup>92</sup> | 2.0.6 |
| edgeR<sup>93</sup> | 4.4.2 |
| limma<sup>94</sup> | 3.62.2 |
| Glimma<sup>95</sup> | 2.16.0 |
| factoextra<sup>96</sup> | 1.0.7 |
| DESeq2<sup>97</sup> | 1.46.0 |
| EnhancedVolcano<sup>98</sup> | 1.24.0 |
| clusterProfiler<sup>99</sup> | 4.14.6 |
| enrichplot<sup>100</sup> | 1.26.6 |
| pheatmap<sup>101</sup> | 1.0.13 |
| FlowJo | 10.10.0 |
| GraphPad Prism | 11.0.2 |

### Cell culture

The CT27 hTSC (XX) line used in the study were derived from first-trimester human placenta CTBs^29^ and were kindly provided by Dr. Hiroaki Okae and Dr. Takahiro Arima (Department of Informative Genetics Environment and Genome Research Center, Tohoku University Graduate School of Medicine, Japan). Chromosomal content of CT27s was validated by Comparative Genomic Hybridization array (Cell Line Genetics). The cells were cultured in human TSC medium (TSCM), composed of a basal medium (DMEM/F12 supplemented with 1% ITS-X, 0.3% BSA, 0.5X penicillin-streptomycin, 0.2 mM L-ascorbic acid) supplemented with 5 μM A83-01, 2 μM CHIR99021, 25 ng/mL EGF, 2.5 μM Y27632, and 0.8 mM VPA. Cells were cultured at 37°C, 5% CO_2,_ and ambient O_2_, and the medium was changed every 2 days. When the TSCs reached 70-80% confluency, they were passaged by detaching with TrypLE diluted 1:1 in 1X PBS for approximately 10 min at 37 °C and were plated onto new dishes with TSCM supplemented with 250 ng/mL iMatrix511.

### Lentivirus production and shRNA-mediated knockdown of *DNMT1*

To generate lentivirus for shRNA KD, HEK293 cells were transfected with either a non-targeting control sequence (shControl) or *DNMT1* targeting pLKO.1 shRNA vectors (shDNMT1_A and shDNMT1_B), pCMV-delta8.9 packaging plasmid, and pCMV-VSV-G envelope plasmid at a molar ratio of 3:2:1 respectively using GenJet reagent, according to the manufacturer’s instructions. After 18 hours, HEK293 medium was replaced with TSCM. Viral supernatants were collected every 24 hours for 2 days. The collected supernatants were filtered through with a SteriFlip Vacuum Tube Top Filter with a 0.45 μm PVDF membrane (Millipore Sigma #SE1M003M00) and frozen at -80 °C.

For lentiviral transduction, hTSC CT27 were plated in 6-well plates at a density of 5x10^4^ cells/well. After 24 hours, the cells were infected with viral supernatants diluted 1:1 with fresh TSCM in the presence of 10 μg/mL polybrene. After an additional 24 hours, the medium was replaced with fresh TSCM. Successfully transduced cells were selected after an additional 24 hours by passaging the cells at a ratio of 1:2 into TSCM supplemented with 250 ng/mL iMatrix511 and 1 μg/mL puromycin. Cells were selected for four days prior to the start of differentiation or maintained in TSCM with 1 μg/mL puromycin for hTSC self-renewal conditions.

### RNA extraction and qPCR

To extract RNA, cells were lysed in TRIzol reagent and total RNA was extracted using phenol-chloroform separation according to the manufacturer’s instructions. cDNA was synthesized from 250-1000ng of RNA using the iScript cDNA synthesis kit. Quantitative PCR (qPCR) was carried out using the PerfeCTa SYBR Green FastMix and 2 μL of cDNA diluted 1:2-1:8 in molecular biology grade H_2_O. The primer sequences for qPCR are provided in Table S1. The data were analyzed using the ΔΔCt method to determine the fold change in expression levels, normalized to glyceraldehyde 3-phosphate dehydrogenase (*GAPDH*) expression.

### Western blot

Cell pellets were lysed in 200 μL 1X RIPA buffer supplemented with cOmplete Protease Inhibitor Cocktail tablets. Protein lysates were denatured using 4X Laemmli Sample Buffer supplemented with 10% β-mercaptoethanol and heated to 95°C. Denatured proteins were separated on 10% SDS-PAGE gels with PageRuler Plus Prestained Protein Ladder as a size standard. After sufficient separation, proteins were transferred onto 0.2 μm PVDF membranes using the Trans-Blot Turbo Transfer System. The membranes were then blocked with 3% BSA in 1X Tris-Buffered Saline with 0.1% Tween 20 (TBST; Cell Signaling Technology #9997). Targets were detected with primary antibodies diluted in 3% BSA-TBST {anti-DNMT1 (1:5000) and anti-GAPDH (1:3000)} followed by incubation with HRP-conjugated secondary antibodies {anti-Mouse-HRP (1:1000) and anti-Rabbit-HRP (1:1000)}. Detection was performed with the ECL Prime Western Blotting System detection reagents. Band intensity was quantified in Bio-Rad Image Lab software and normalized to GAPDH as a loading control.

### Immunofluorescent staining

For all targets apart from HLA-G, cells grown in black-walled 96-well plates were fixed with 4% PFA followed by permeabilization with 1X PBS with 0.4% Triton X-100 (PBS-Triton). Cells were then blocked with 3% BSA in PBS. Targets were detected with primary antibodies diluted in 3% BSA {anti-TFAP2C (1:500) and anti-KRT7 (1:500)} followed by fluorescent labeled secondary antibodies {anti-Rabbit-AF555 (1:1000) and anti-Mouse-AF647 (1:1000)}. Finally, cells were stained with 10 μg/mL DAPI in TBS for 5 minutes. Images were acquired on a Nikon ECLIPSE Ti-2 inverted widefield microscope. Fluorescent intensity was quantified in nuclei or cytoplasmic areas, depending on protein localization, in FIJI^83^.

For HLA-G, cells grown in black-walled 96-well plates were stained live with PE-conjugated HLA-G antibody (1:300) diluted in cell culture medium for 15 minutes at 37 °C. Cells were then fixed in 4% PFA, permeabilized, stained with DAPI, and imaged as described above.

### Cell proliferation assay

To assess cell proliferation in hTSCs, cells were plated at normalized cell densities in black-walled 96-well plates at day 12 post-transduction. After 24 hours, at day 13 post-transduction, 10 μM EdU was added to the medium and cells were incubated for one hour at 37 °C. Cells were then fixed with 4% PFA for 15 minutes and permeabilized with PBS-Triton for 20 minutes. EdU was detected with the Click-iT Plus EdU Alexa Fluor 488 dye kit prepared according to manufacturer’s instructions. Cells were stained with DAPI and imaged as described for immunofluorescent assays. The portion of nuclei positive for EdU was determined in FIJI^83^.

### Low-depth whole genome bisulfite sequencing (Sparse-Seq) and analysis

At day 13 post-transduction, shRNA transduced hTSCs were collected in cell pellets and frozen at -80C. Pellets were then thawed on ice, and DNA was extracted using the QIAamp DNA Mini Kit. Residual RNA was removed by treatment with 10 μg/mL RNAse at 37 °C for 5 minutes, followed by clean-up of genomic DNA with the QIAamp DNA Micro Kit. 200ng of purified DNA per sample was then subjected to bisulfite conversion with the EpiTect Bisulfite Kit, followed by secondary strand synthesis, tagmentation, and library amplification and purification using the Zymo-Seq WGBS Library Kit and the Illumina Nextera DNA Flex Library Prep Kit. Library size was analyzed by Agilent TapeStation D1000 ScreenTape Assay and quantified by qPCR using the KAPA Library Quantification Kit. Libraries were then sequenced on an Illumina MiSeq i100 with 300bp paired end reads at a depth of 500,000 to 1 million reads per sample.

To analyze Sparse-Seq data, reads were trimmed of adaptor sequences and low-quality bases using the trim-galore package^84^. Sequencing quality was assessed by FastQC^85^. Reads were then aligned and methylation values extracted using Bismark^86^. Extracted methylation values were quantified and plotted in R in RStudio using ggplot2^87^. Genomic regions and repeat elements were defined using GenomicRanges^88^ and rtracklayer^89^, respectively.

### ChromHMM analysis

Publicly available ChIP-seq and CUT&TAG data was downloaded from the Gene Expression Omnibus (GEO) database (Table S2). For analysis, reads were first trimmed of adaptor sequences and low-quality bases using the trim-galore package^84^. Trimmed reads were then aligned to the hg38 human genome using HISAT2^90^ and resulting .sam files were sorted and converted to .bam files using SAMtools^91^. ChromHMM^39,40^ analysis was then performed with state 12 emissions. Methylation measured by Sparse-Seq at ChromHMM defined states was then plotted in R in RStudio using ggplot2^87^.

### Differentiation of hTSCs into hTSC-EVTs and hTSC-STs

Differentiation of hTSC CT27 into hTSC-EVTs was performed using a previously described protocol^29^. Briefly, hTSCs were dissociated with TrypLE diluted 1:1 in PBS then plated onto 1 μg/mL collagen IV-coated dishes at a density of 1-1.5x10^4^ cells/cm^2^ with 2 mL of EVT differentiation medium, which contained DMEM/F12 medium supplemented with 1% ITS-X, 0.5% penicillin-streptomycin, 0.3% BSA, 0.1 mM β-mercaptoethanol, 100 ng/mL NRG1, 7.5 μM A83-01, 2.5 μM Y27632, 4% KSR, and 2% Matrigel supplemented with 1 μg/mL puromycin for lentiviral based KD experiments. On day 3, the EVT medium was replaced with fresh medium supplemented with reduced Matrigel (0.5%) and without NRG1. On day 6, the EVT medium was again replaced with medium containing reduced Matrigel (0.5%) and without both NRG1 and KSR. hTSC-EVTs were analyzed after eight days of differentiation.

Differentiation of hTSC CT27 into hTSC-STs was performed using a previously described 3D protocol^29^. Briefly, hTSCs were dissociated using TrypLE diluted 1:1 in PBS and then cultured in low attachment 6-well plates at a density of 1-2.5x10^5^ cells/well with 2 mL of ST(3D) differentiation medium supplemented with 1 μg/mL puromycin. The ST(3D) differentiation medium was composed of DMEM/F12 supplemented with 1% ITS-X supplement, 0.5% penicillin-streptomycin, 0.3% BSA, 0.1 mM β-mercaptoethanol, 2.5 μM Y27632, 2 μM forskolin, 50 ng/mL EGF, and 4% KSR. The medium was replaced after three days, and hTSC-STs were analyzed after six days of differentiation.

### Flow cytometry

hTSC-EVTs were assessed for HLA-G positivity by flow cytometry after eight days of differentiation. Cells were dissociated using TrypLE and quenched with FACS buffer (PBS supplemented with 1% BSA and 5mM EDTA). Cells were stained live with anti-HLA-G-PE diluted at 1:100 in FACS buffer for one hour on ice. Dead cells were labelled with a LIVE/DEAD Fixable Violet Dead Cell Stain diluted at 1:1000 in PBS for 30 minutes on ice. Finally, cells were fixed in 2% PFA in FACS buffer for 10 minutes on ice. Flow cytometry was performed on a BD LSRFortessa. Samples were compared to an unstained negative control, and analysis was carried out in FlowJo software.

### Enzyme-Linked Immunosorbent Assays (ELISA)

Production of β-HCG by hTSC-STs was assayed by ELISA. Medium was exchanged on day 3 of differentiation, following the standard differentiation protocol, and collected on day 6 of differentiation (day 12 post-transduction). β-HCG content of this medium was assessed by clinical ELISA performed by the Andrology Laboratory at the University of Pennsylvania Health System Fertility Care clinic.

Production of CXCL2 in hTSCs and hTSC-EVTs was assayed by ELISA. For undifferentiated hTSCs, cells were seeded at normalized cell density on day 11 post-transduction and conditioned medium was collected 48 hours later (day 13 post-transduction) and flash frozen. For hTSC-EVTs medium was replaced on day 6 of differentiation, following the standard protocol, and then collected 48 hours later at day 8 of differentiation (day 14 post-transduction) and flash frozen. Samples were thawed on ice prior to ELISA protocol following manufacturer’s instructions.

### Bulk RNA sequencing and analysis

Cell pellets for RNA sequencing were collected at day 13 post-transduction for the undifferentiated hTSC condition and after eight days of differentiation (14 days post-transduction) for the differentiated EVT condition and frozen at -80°C. RNA extraction, library preparation, and sequencing was performed by Azenta Life Sciences. Libraries were sequenced on an Illumina NovaSeq with 150bp paired end reads at a depth of 25-30 million reads per sample.

For analysis, .fastq files were processed to .bam files in the same manner as for ChromHMM analysis. Reads were then summarized using the featureCounts^102^ function in the Rsubread package^92^. Read counts were then filtered to remove genes with a read count of zero for all samples and normalized using edgeR^93^, limma^94^, and Glimma^95^. Principal component analysis (PCA) was performed using factoextra^96^. Differential gene expression analysis was performed using DESeq2^97^ with thresholds of |log_2_ fold change| > 1 and adjusted p value < 0.05. Gene set enrichment analysis (GSEA), functional enrichment, visualization of differential gene expression analysis were performed using EnhancedVolcano^98^, clusterProfiler^99^, and pheatmap^101^.

### Treatment with DNMT1 inhibitors

To determine if the role of DNMT1 in EVT differentiation depends on its catalytic function as a DNA methyltransferase, we treated hTSCs with a catalytic DNMT1 inhibitor^64^ (DNMT1i; GSK3685032). hTSC CT27 were seeded at a density of 2.1x10^3^ cells/cm^2^ in hTSC media with iMatrix511. After 24 hours, media was exchanged for hTSC media supplemented 10 μM DNMT1i or DMSO as a vehicle control. After 48 hours, medium was exchanged with fresh medium, supplemented with DNMT1i or DMSO. After 72 hours of treatment, cells were differentiated into EVTs, with continued treatment with DNMT1i or DMSO during differentiation. After the eight-day EVT differentiation protocol, inhibitor function and EVT differentiation efficiency were assessed.

### Targeted methylation sequencing and analysis

At day 8 of EVT differentiation with DNMT1i treatment, cells were collected in pellets and frozen at -80C. Pellets were thawed on ice, and DNA was extracted using the QIAamp DNA Mini Kit. 1000 ng of purified DNA per sample bisulfite converted using the EpiTect Bisulfite Kit. The targeted region of interest was amplified with the PyroMark PCR kit using primers designed using BiSearch^103^ primer design tool with overhangs for complementarity to index primers (Table S1). Amplification of the *KCNQ1OT1*:TSS-DMR target region (chr11:2699149-2699412) was validated by agarose gel electrophoresis. Indexing of amplicons for Illumina sequencing was performed using the Multiplex PCR Kit. Indexed libraries were then purified using AMPure XP Beads. Validation of library size, quantification, and sequencing was performed as described in Sparse-Seq methods with greater than 1000 reads per amplicon. Read trimming, QC, alignment, and methylation extraction were performed as described in Sparse-Seq methods. Methylation values were averaged across all CpGs per locus.

### Invasion Assay

EVT invasion was assessed in a modified invasion assay protocol^48,104^. Briefly, SecureSeal Hybridization Chambers (Grace BioLabs #621201) chambers were sterilized with ultraviolet light in a biosafety cabinet for one hour. Chambers were then adhered to the bottom of 12 well plates. EVT medium was mixed with Matrigel at a ratio of 1:2 and 60 μL of this mixture was loaded into each hybridization chamber. The Matrigel-EVT media mixture was allowed to solidify for one hour at 37°C. DNMT1i or DMSO treated hTSCs were diluted in EVT medium without Matrigel at a concentration of 2x10^5^ cells/mL and 50 μL of cells (1x10^4^ cells) were loaded in each chamber. Plates were incubated upright for one hour at 37°C to allow cells to settle on the edge of the solidified Matrigel-EVT media mixture. After this time, 1.5mL EVT media without Matrigel with either 10μM DNMT1i or DMSO vehicle control was added to each well. Medium was then exchanged on days 3 and 6 of differentiation. Images to measure invasion depth were captured on an EVOS M7000 Imaging System on day 8 of EVT differentiation. Depth of invasion was measured in six areas per replicate in FIJI^83^ and averaged.

### Statistical analysis

All statistical tests used are indicated in the figure legends. Statistical tests were performed in GraphPad Prism, except for tests comparing methylation values in Figure 2 and expression of the EVT gene set in Figure 4, which were performed in R. All error bars represent the standard error of the mean.

### Data availability

The RNA-sequencing and low-depth, bisulfite-sequencing data generated in this paper are available in the Gene Expression Omnibus (GEO: GSE337276 & GSE337277). This study analyzed publicly available data detailed in Table S2. This paper does not report any original code.

## Acknowledgements

We would like to acknowledge Dr. Hiroaki Okae and Dr. Takahiro Arima (Tohoku University) for kindly providing the hTSC cell line used in this study. We would also like to acknowledge Dr. Marisa Bartolomei and members of her lab for their guidance in this project and reading of the manuscript.

## Author Contributions

Conceptualization, W.M., J.M.K., J.K.; Investigation, S.L.K, Y.J.J., G.G., E.D.T, J.C.; Analysis, S.L.K, Y.J.J., G.G., L.S.P.; Writing – Original Manuscript, S.L.K, Y.J.J., G.G.; Writing – Review and Editing, S.L.K, Y.J.J., G.G., E.D.T, L.S.P., J.C., J.K., W.M., J.M.K

