## Supplemental Figures and Tables for "DNA methylation maintenance by DNMT1 is essential for human trophoblast stem cell homeostasis and differentiation"

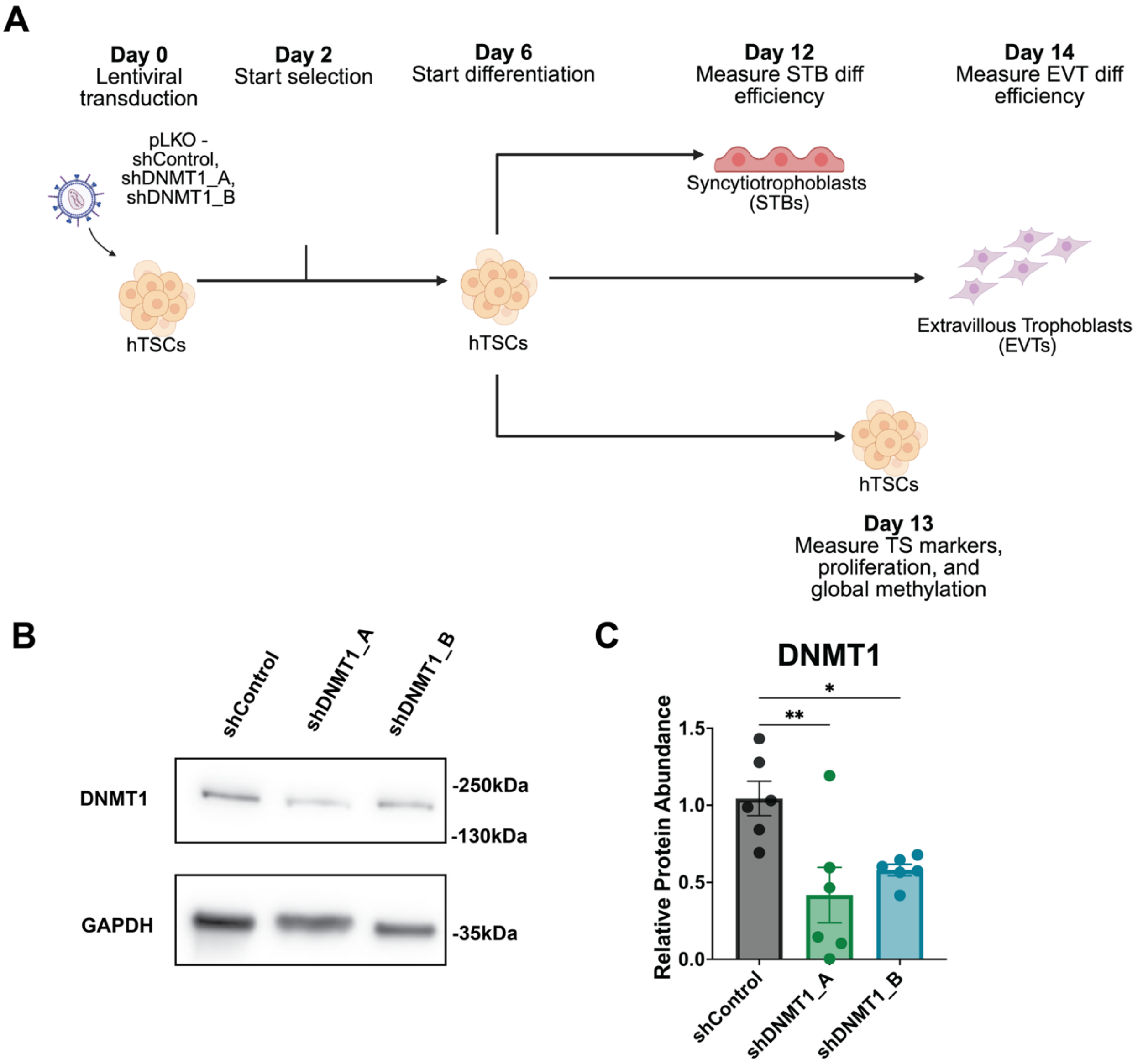


**Supplemental Figure 1: Representation and validation of *DNMT1* KD in hTSCs.** A) Schematic representation of *DNMT1* KD experiments in hTSC condition and ST and EVT differentiation conditions. (Created in BioRender. Tichy, E. (2026) https://BioRender.com/xa8xk7y) B) Representative western blot image to validate DNMT1 KD efficiency at day 13 post-transduction C) Quantification of western blots show decreased DNMT1 protein abundance at day 13 post-transduction normalized to GAPDH (One-way ANOVA with multiple comparisons vs. shControl, * = $p < 0.05$, ** = $p < 0.01$, n = 6 biological replicates).

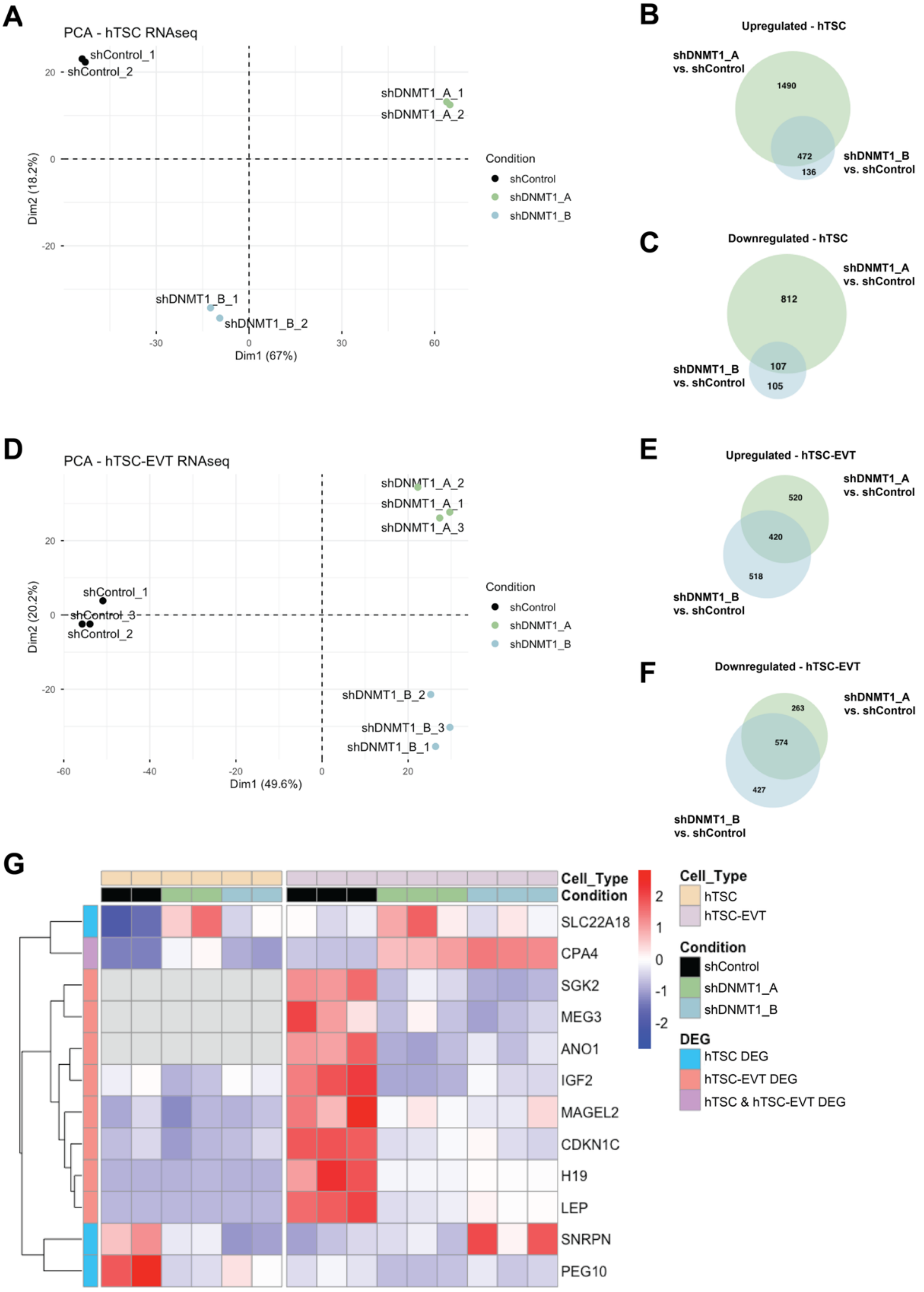

A
PCA - hTSC RNAseq
shControl_1
shControl_2
shDNMT1_A_1
shDNMT1_A_2
shDNMT1_B_1
shDNMT1_B_2
Dim1 (67%)
Dim2 (18.2%)
Condition
shControl
shDNMT1_A
shDNMT1_B
B
Upregulated - hTSC
shDNMT1_A vs. shControl
shDNMT1_B vs. shControl
1490
472
136
C
Downregulated - hTSC
shDNMT1_A vs. shControl
shDNMT1_B vs. shControl
812
107
105
D
PCA - hTSC-EVT RNAseq
shDNMT1_A_2
shDNMT1_A_1
shDNMT1_A_3
shControl_1
shControl_3
shControl_2
shDNMT1_B_2
shDNMT1_B_3
shDNMT1_B_1
Dim1 (49.6%)
Dim2 (20.2%)
Condition
shControl
shDNMT1_A
shDNMT1_B
E
Upregulated - hTSC-EVT
shDNMT1_A vs. shControl
shDNMT1_B vs. shControl
520
420
518
F
Downregulated - hTSC-EVT
shDNMT1_A vs. shControl
shDNMT1_B vs. shControl
263
574
427
G
Cell_Type
Condition
SLC22A18
CPA4
SGK2
MEG3
ANO1
IGF2
MAGEL2
CDKN1C
H19
LEP
SNRPN
PEG10
Cell_Type
hTSC
hTSC-EVT
Condition
shControl
shDNMT1_A
shDNMT1_B
DEG
hTSC DEG
hTSC-EVT DEG
hTSC & hTSC-EVT DEG

**Supplemental Figure 2: *DNMT1* KD impacts the hTSC-EVT transcriptome.** A) Principal components analysis of hTSC RNA-seq for shControl, shDNMT1_A, and shDNMT1_B shows clear clustering of replicates. Venn diagrams of B) upregulated and C) downregulated genes in hTSCs demonstrate overlap in shDNMT1_A and shDNMT1_B vs shControl comparisons. D) Principal components analysis of hTSC-EVT RNA-seq for shControl, shDNMT1_A, and shDNMT1_B shows clear clustering of replicates and separation of *DNMT1* KD conditions from controls on Dim1. Venn diagrams of E) upregulated and F) downregulated genes in hTSC-EVTs demonstrate overlap in shDNMT1_A and shDNMT1_B vs shControl comparisons. Overlapping up- or downregulated genes were used for all subsequent analyses. G) Heatmap of differentially expressed imprinted genes in *DNMT1* KD hTSCs and hTSC-EVTs. (hTSC DEG – differentially expressed in *DNMT1* KD hTSCs, hTSC-EVT DEG – differentially expressed in *DNMT1* KD hTSC-EVTs, hTSC & hTSC-EVT DEG – differentially expressed in *DNMT1* KD hTSCs and hTSC-EVTs).

**Supplemental Table 1: Oligonucleotide sequences for qPCR and targeted bisulfite sequencing**

| Primer | Forward Primer | Reverse Primer |
|---|---|---|
| *GAPDH*_qPCR | TGACTTCAACAGCGACACCC | TTCGTTGTCATACCAGGAAATGAG |
| *DNMT1*_qPCR | GACAGTGATGAGCAGCCCAT | CCTGTCCTTCTCCCTGGTA |
| *GATA3*_qPCR | GGGCTCTACTACAAGCTTCACAATA | CACTTTTTGGATTTGCTAGACATTT |
| *TEAD4*_qPCR | CAGGTGGTGGAGAAAGTTGAGA | GTGCTTGAGCTTGTGGATGAAG |
| *TP63*_qPCR | AGAAACGAAGATCCCCAGATGA | CTGTTGCTGTTGCCTGTACGTT |
| *HLA-G*_qPCR | CCACCACCCTGTCTTTGACTAT | ACGTCCTGGGTCTGGTCCT |
| *MMP2*_qPCR | TGGCACCCATTTACACCTACAC | ATGTCAGGAGAGGCCCCATAGA |
| *ASCL2*_qPCR | GCACCAACACTTGGAGATTTT | AATGGATTCTCTGTGCCCTTAG |
| *CGB*_qPCR | CAGCATCCTATCACCTCCTGGT | CTGGAACATCTCCATCCTTGGT |
| *SDC1*_qPCR | CTATTCCCACGTCTCCAGAACC | GGACTACAGCCTCTCCCTCCTT |
| *ERVW-1*_qPCR | CGAACGGACATCCAAAGTG | GACTCAAGTGTGATGTATCCAAGACT |
| *KCNQ1OT1*:TSS-DMR_BIS | ACACTCTTTCCCTACACGACGCTCTTCCGATCTTTTATTTTGAATTATTATGAGAA | GTGACTGGAGTTCAGACGTGTGCTCTTCCGATCTTCCCCATCTCTCTAAAAAAA |

**Supplemental Table 2: Data for ChromHMM Analysis**

| GEO Accession Number | Sample Number | Chromatin Mark | Method | Cell Line | DOI |
|---|---|---|---|---|---|
| GSE200759 | GSM6043300 | H3K27ac | ChIP-seq | CT27 | 10.1038/s41594-023-00960-6[1] |
| GSE204722 | GSM7498141 | H3K27ac | ChIP-seq | CT27 | 10.1038/s41467-023-40424-5[2] |
| GSE266194 | GSM8242200 | H3K27me3 | ChIP-seq | CT29 | 10.1016/j.stem.2024.12.007[3] |
| GSE266194 | GSM8242192 | H3K27me3 | ChIP-seq | BTS11 | 10.1016/j.stem.2024.12.007[3] |
| GSE196221 | GSM5862947 | H3K36me3 | ChIP-seq | Zhou-hTSC | 10.1073/pnas.2216206120[4] |
| GSE266194 | GSM8242190 | H3K36me3 | ChIP-seq | BTS11 | 10.1016/j.stem.2024.12.007[3] |
| GSE200759 | GSM6043301 | H3K4me1 | ChIP-seq | CT27 | 10.1038/s41594-023-00960-6[1] |
| GSE200761 | GSM6043315 | H3K4me1 | CUT&TAG | CT27 | 10.1038/s41594-023-00960-6[1] |
| GSE196221 | GSM5862950 | H3K4me3 | ChIP-seq | Zhou-hTSC | 10.1073/pnas.2216206120[4] |
| GSE204722 | GSM7498142 | H3K4me3 | ChIP-seq | CT27 | 10.1038/s41467-023-40424-5[2] |
| GSE266194 | GSM8242201 | H3K9me3 | ChIP-seq | CT29 | 10.1016/j.stem.2024.12.007[3] |
| GSE266194 | GSM8242193 | H3K9me3 | ChIP-seq | BTS11 | 10.1016/j.stem.2024.12.007[3] |
| GSE196221 | GSM5862951 | Input | ChIP-seq | Zhou-hTSC | 10.1073/pnas.2216206120[4] |
| GSE200759 | GSM6043303 | Input | ChIP-seq | CT27 | 10.1038/s41594-023-00960-6[1] |
| GSE200761 | GSM6043319 | Input | CUT&TAG | CT27 | 10.1038/s41594-023-00960-6[1] |
| GSE204722 | GSM7498140 | Input | ChIP-seq | CT27 | 10.1038/s41467-023-40424-5[2] |
| GSE266194 | GSM8242194 | Input | ChIP-seq | BTS11 | 10.1016/j.stem.2024.12.007[3] |
